# Prion propagation is dependent on key amino acids in Charge cluster 2 within the prion protein

**DOI:** 10.1101/2022.08.08.503133

**Authors:** Savroop Bhamra, Parineeta Arora, Szymon W. Manka, Christian Schmidt, Craig Brown, Melissa L. D. Rayner, Peter-Christian Klöhn, Anthony R. Clarke, John Collinge, Parmjit S. Jat

## Abstract

Prions consist of assemblies of aberrantly folded cellular prion protein (PrP^C^) upon template-assisted conversion and propagation of disease-associated PrP. To dissect the N-terminal residues critical for efficient prion propagation, we generated a library of point, double, or triple alanine replacements within residues 23-111 of PrP, stably expressed them in cells silenced for endogenous mouse PrP^C^ and challenged the reconstituted cells with four mouse prion strains. Amino acids (aa) 105-111 of Charge Cluster 2 (CC2), which is disordered in PrP^C^, were required for propagation of all four prion strains; other residues had no effect or exhibited strain-specific effects. Replacements in CC2, including aa105-111, dominantly inhibited prion propagation in the presence of endogenous wild type PrP^C^ whilst other changes were not inhibitory. Single alanine replacements within aa105-111 identified leucine 108 and valine 111 or the cluster of lysine 105, threonine 106 and asparagine 107 as critical for prion propagation. These residues mediate specific ordering of CC2 in the prion fibrils from Rocky Mountain Laboratory (RML) and ME7 mouse prion strains.

## Introduction

Prion diseases are fatal progressive neurodegenerative maladies that involve accumulation of assemblies of an aberrantly folded PrP^C^. Misfolding of PrP^C^ is an autocatalytic process of seeded fibrillization and fission and involves dramatic changes to the conformation of PrP, from α-helix-rich to a β-sheet-rich protein (Prusiner, 1998) (Collinge & Clarke, 2007) (Collinge, 2016) (Terry & Wadsworth, 2019). Recent electron microscopic analyses of high-titre exceptionally pure infectious prions have identified rods comprised of single-protofilament helical amyloid fibrils (Kraus *et al*,2021) (Manka *et al*, 2022b) and twisted pairs of the same protofilaments (Terry *et al*, 2016) (Terry *et al*, 2019) (Manka *et al*., 2022b) (Manka *et al*, 2022a); each rung of the protofilament is a single PrP monomer.

PrP^C^ is a highly conserved cell-surface glycoprotein, expressed in most cell types, but without a precise unitary cellular function, although many roles have been proposed (Linden *et al*, 2008) (Linden *et al*, 2011) (Linden, 2017). The wide variety of activities and functions ascribed to PrP^C^ suggest that it does not act exclusively in a single pathway, but may function as a dynamic scaffold for the assembly of various multicomponent signalling complexes while it moves between the cell surface and the endocytic compartment (Linden *et al*., 2008) (Linden *et al*, 2011) (Linden, 2017). PrP^C^ undergoes autocatalytic fibrillization and fission after coming into contact with prions at the cell surface (Goold *et al*, 2011), with the newly formed prions residing at the cell surface for hours in string-and web-like structures attached to the cell membrane (Rouvinski *et al*, 2014).

Mice devoid of PrP (*Prnp^0/0^*) are viable, have no overt phenotype but are completely resistant to prion disease indicating that PrP is essential for pathogenesis (Bueler *et al*, 1992) (Bueler *et al*, 1993) (Sakaguchi *et al*, 1995) (Fischer *et al*, 1996). Development of these *Prnp^0/0^* mice led to an extensive structure-function analysis of PrP to identify domains that are important for prion propagation by reconstituting these mice with mutant PrP transgenes. This demonstrated that N-terminally truncated PrP (Δ23-80, Δ32-80 and Δ32-93) could be converted to infectious prions but at a reduced level of susceptibility (Fischer *et al*., 1996; Flechsig *et al*, 2000), whereas PrP Δ32-106 could not be converted to disease-associated PrP nor supported RML prion propagation upon reconstitution of *Prnp^0/0^* mice (Weissmann & Flechsig, 2003). This finding that the N-terminus was essential for efficient prion propagation was further supported by Suppatapone *et al* who found that reconstitution of *Prnp^0/0^* mice with PrP Δ23-88 propagated prions but with a longer incubation time (Supattapone *et al*, 2001).

The N-terminal half of PrP^C^ is glycine-rich and disordered. It encompasses two charge clusters, denoted CC1 (polybasic region, 23.KKRPK.27) and CC2 (90.QGGGTHNQWNKPSKPKTNLKHV.111), octapeptide repeats (OPRs), and an alanine-rich low-complexity region (LCR, 112.AGAAAAGAVVGGLGG.126). The CC1 domain is fundamental for: (i) PrP^C^ endocytosis via coated pits (Taylor *et al*, 2005); (ii) overall folding of the globular C-terminal domain (Ostapchenko *et al*, 2008); (iii) efficiency of prion propagation in transgenic mice (Turnbaugh *et al*, 2012). The octapeptide repeat region (OPR, 51-90), contains one cryptic and four consensus repeats of a series of eight amino acids. Supernumerary insertions of between one and nine additional OPRs may increase the risk of developing disease with most cases showing an earlier onset (Stevens *et al*, 2009). Deletion mutagenesis studies have suggested that the OPRs are not required for prion propagation, and play only a limited role in disease pathogenesis (Yamaguchi *et al*, 2012). The function of OPRs remains unclear but they can bind divalent metal ions through coordination to histidine residues and may play a role in copper homeostasis (Jackson *et al*, 2001). CC2 is one of the most immunogenic regions of PrP^C^ and reported to be the interaction site for amyloid-beta oligomers (Lauren *et al*, 2009) (Chen *et al*, 2010) (Freir *et al*, 2011) (Fluharty *et al*, 2013) (Rushworth *et al*, 2013), heparin and copper (Warner *et al*, 2002). It is also proposed to specifically bind disease-associated PrP, leading to seeded misfolding of PrP^C^ (Abalos *et al*, 2008).

In addition to reconstitution of *Prnp^0/0^* mice, many of these mutant transgenes were also used to study prion propagation *in vitro* in cells expressing endogenous PrP^C^, or chronically prion-infected N2a (ScN2a) cells (Rogers *et al,* 1993) (Priola *et al,* 1994) (Abalos *et al,* 2008) (Shirai *et al,* 2014). However these results may have been compromised by the presence of endogenous PrP as Supattapone *et al*. observed that upon RML infection, mouse wild-type PrP (moPrP^WT^) can act in *trans* to accelerate propagation of a PrP double deletion mutant (Δ23-88 and Δ141-176) in transgenic mice (Supattapone *et al*, 1999). There is a further complication in that many previous studies have used the 3F4 epitope to distinguish the exogenously introduced protein from the endogenous protein. The epitope for the 3F4 antibody is generated by mutating leucine 108 and valine 111 to methionine in mouse PrP. Suppatapone *et al* found that in transgenic mice, the presence of the 3F4 epitope in PrP has a non-strain specific adverse effect on prion propagation (Supattapone *et al*., 2001).

Just as the development of *Prnp*^0/0^ mice paved the way for a structure-function analysis of PrP in mice by transgenic expression of N-terminal deletion mutants (Bueler *et al*., 1992) (Fischer *et al*., 1996), our aim was to identify which residues within the unstructured amino-terminal domain were required for prion propagation in cells. We determined this by: (i) generating cells stably knocked-down for PrP^C^ expression to a level that renders them fully resistant to prion infection, while regaining susceptibility to infection upon restoring PrP expression; (ii) preparing a library of single, double and triple replacements of all residues, except proline and glycine, to alanine within the N-terminal unstructured 23-111 region and assaying their ability to support propagation of RML prions. We extended our study to three other prion strains, ME7, 22L and MRC2 in CAD5 cells. Like RML, 22L and ME7 are mouse-adapted scrapie prion strains with a relatively short incubation period, whereas MRC2 is a mouse-adapted bovine spongiform encephalopathy (BSE) prion strain characterised by a long incubation time and di-glycosylated-dominant PrP^Sc^ (Lloyd *et al*, 2004a). The rationale for this mutagenesis approach was informed by extensive research showing that elimination of sidechains beyond the β-carbon by alanine replacement, with minimal perturbation of the protein backbone conformation, can probe the influence of specific amino acid side-chains on biological activity, protein stability or folding (Cunningham & Wells, 1989) (Fersht & Daggett, 2002). It is applicable to a wide range of amino acids, as it contains an inert, non-bulky methyl group, and retains the secondary structure preferences of many amino acids, thus minimally affecting secondary structure (Fersht & Daggett, 2002) (Tang & Fenton, 2017). Proline and glycine residues were not targetted for mutagenesis in this study to minimise the amount of structural change introduced in the protein as they provide rigidity and flexibility respectively, to the protein backbone. We show that efficient prion propagation is strain dependent and contingent upon leucine 108 and valine 111 acting alone or lysine 105, threonine 106 and asparagine 107, acting together at the infection stage.

## Results

### PK1-KD cells: a stable cell line knocked-down for expression of endogenous prion protein

To study the effect of *Prnp* mutations on prion propagation without interference from endogenous PrP^C^, we stably silenced it in PK1-10 cells, a cell line highly susceptible to RML infection, using the pRetroSuper vector system. PK1-10, are a single cell clone of the PK1 murine neuroblastoma cell line which is a clone of mouse N2a neuroblastoma cells that are highly susceptible to prion infection and able to maintain a chronic RML prion infection (Klohn *et al*, 2003). PK1-10 cells are referred to as PK1 from hereon as the only sub-clone of PK1 cells used in this study.

Eight different hairpins, designed to predominantly target the 3’ untranslated region (UTR) of endogenous PrP^C^ (Fig.1A), were stably expressed in PK1 cells. Bulk cultures of stably transduced cells were prepared and western blotted to determine PrP^C^ expression; shRNA8 markedly reduced PrP expression. Retrovirus encoding shRNA8 was used to repeat transduction of PK1 cells and isolate single cell clones. 96 clones were isolated, reconstituted with the full-length open reading frame (ORF) for moPrP^WT^ (pLNCX2moPrP^WT^) and challenged with RML prions in a scrapie cell assay (SCA), a highly sensitive quantitative cell-based infectivity assay (Klohn *et al*., 2003). In SCA, a pre-determined number of cultured cells are infected with prions and serially passaged for three splits to dilute out the original innoculum. At the 4^th^, 5^th^ and 6^th^ splits, the number of cells containing PK-resistant PrP (PrP^Sc^) is assessed via ELISPOT using the anti-PrP antibody, ICSM18. Cells containing PK-resistant PrP at this stage represent stably infected cells propagating prions. The relationship between spot number and infectivity as measured in Tissue culture infectious units (TCIU) is not linear (detailed in Fig. EV1). Split 6 data is presented within the body of the paper while complete data sets (4^th^, 5^th^ and 6^th^ splits) are provided as extended data.

**Figure 1.**
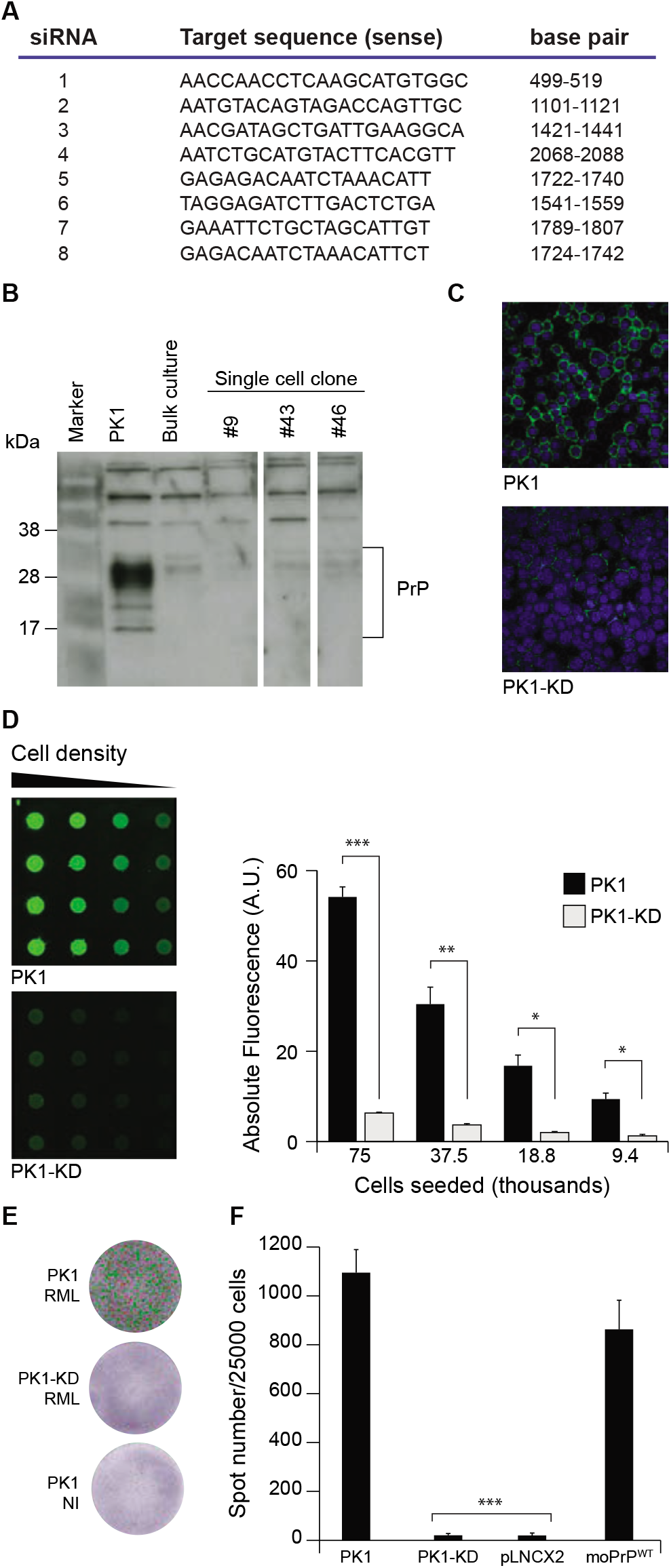
PK1-KD cells. A. Eight siRNA constructs were designed against the native mouse PrP sequence and used to develop shRNA constructs for stable silencing of expression. Of these, shRNA8 was used to establish cells knocked-down for mouse PrP expression. B. Western blot showing PrP detected in PK1 cells; loss of PrP expression was observed in the bulk culture where shRNA8 was used for silencing. Higher level of suppression of PrP expression was observed in clones 9, 43 and 46. Clone 9 was selected for this study. PK1shRNA8 clone 9 was designated as PK1-KD cells. C. Immunofluorescence data showing PrP expression in PK1 and PK1-KD cells respectively. DAPI, nuclear stain; mouse PrP, green. D. Quantification of protein expression by dot blot showed that the level of PrP silencing achieved in PK1siRNA8 clone 9 was 88%. *P-values* were calculated using an unpaired t-test; *** denotes *p*≤0.0001, ** *p*≤0.0005 and * *p*≤ 0.005. E. Images of individual ELISPOT wells showing one with typically high spot numbers for PK1 cells infected with RML (1000 spots) and low spot numbers for both RML infected PK1-KD cells and non-infected (NI) PK1 cells (23 and 10 spots respectively). Positive (green) and negative (red) spots are highlighted by the imaging software over a purple background from colour development in the ELISPOT procedure. F. SCA data showing spot numbers detected upon infection with RML [10^-5^ dilution of 10% infected brain homogenate (BH)]. PK1 cells and PK1-KD cells reconstituted with moPrP^WT^ report full susceptibility to infection (>500 spots), while PK1-KD cells and those transduced with the empty vector, pLNCX2, were resistant to RML infection (<50 spots). Significance was calculated in a one-way ANOVA with Bonferroni correction for multiple comparisons to PK1-KD cells reconstituted with moPrP^WT^. It is indicated by *** for *p*≤0.0001.

Three clones isolated from PK1shRNA8 bulk culture (9, 43 and 46) that did not propagate prions but regained full susceptibility to RML upon reconstitution with moPrP^WT^, indicated by increasing numbers of PrP^Sc^-containing cells upon passaging, were expanded and frozen. PK1shRNA8 clone 9 was one such clone (Fig. 1B and C); it has been designated PK1-KD and was used for the reconstitution experiments. Compared with non-silenced cells, the level of endogenous PrP in PK1-KD cells was reduced by 88% as determined by dot blot (Fig. 1D). PK1-KD cells are resistant to RML infection (Fig. 1E), but when reconstituted with moPrP^WT^ regain full susceptibility to infection (Fig. 1F).

### Propagation of RML prions is modulated by three distinct domains in the N-terminus of PrP

Suppatapone *et al* found that N-terminal amino acids 23-88 (Fig. 2A) were required in transgenic mice for efficient prion propagation. To determine if this would be replicated in cultured cells, we reconstituted PK1-KD cells with moPrP Δ23-88 (Fig.2B) and challenged the resulting bulk cultures with RML prions (Fig. 2C and Fig. EV2A). The effect of Δ23-88 was stronger in PK-1KD cells than in *Prnp*^0/0^ mice, as it completely abrogated prion propagation. To determine which specific N-terminal residues were critical for prion propagation, single, double and triple replacements to alanine were prepared (Fig. 3A) and used to reconstitute PK1-KD cells and bulk cultures challenged with RML (Fig. 3B and D). Two sites, residues 23-25 (KKR), within CC1 domain and glutamine 41 were found to be required. Mutants K23A.K24A.R25A and Q41A exhibited considerably reduced RML propagation (Fig. 3B and Fig. EV2B). While cells that fully supported prion propagation consistently reported over 800 PrP^Sc^ containing cells (spot number), less than 200 were observed for both K23A.K24A.R25A and Q41A mutants. This spot number is much lower compared with PK1 and PK1-KDmoPrP^WT^ cells, yet greater than PK1-KD and PK1-KDΔ23-88 cells (which do not propagate RML prions) suggesting that prion propagation is markedly compromised, but not abrogated in mutants K23A.K24A.R25A and Q41A.

**Figure 2.**
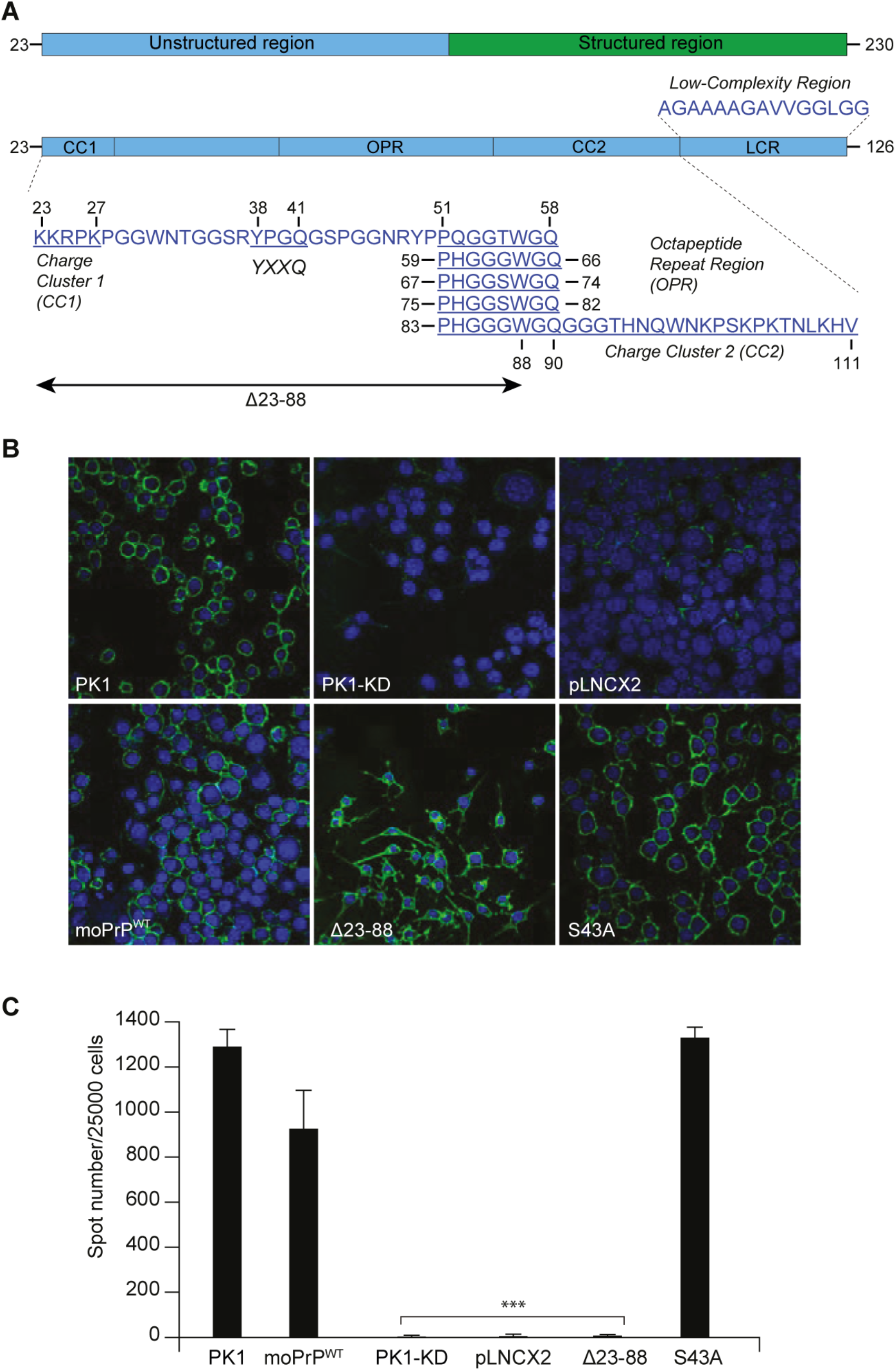
N-terminal sites in moPrP targeted for mutagenesis. A. Schematic showing the full-length mature mouse prion protein (moPrP, aa23-230) with regions of the unstructured amino-terminal domain highlighted as CC1 (Charge Cluster 1), OPR (Octapeptide Repeats), CC2 (Charge Cluster 2) and LCR (Low-Complexity Region). The amino acid sequence is displayed below or above the bars in single letter code with regions of interest highlighted (mouse PrP numbering). B. Immunofluorescence images showing PrP expression in PK1, PK1-KD and PK1-KD cells reconstituted with empty vector pLNCX2, moPrP^WT^, mutant Δ23-88 and the alanine mutant S43A. DAPI, nuclear stain; mouse PrP, green. C. SCA data showing spot numbers for PK1, PK1-KD and reconstituted PK1-KD cells at split 6, post-RML infection [10^-5^ dilution of 10% infected brain homogenate]. For SCAs, significance was calculated in a one-way ANOVA with Bonferroni correction for multiple comparisons to PK1-KD cells reconstituted with moPrP^WT^. It is indicated by *** for *p*≤0.0001.

**Figure 3.**
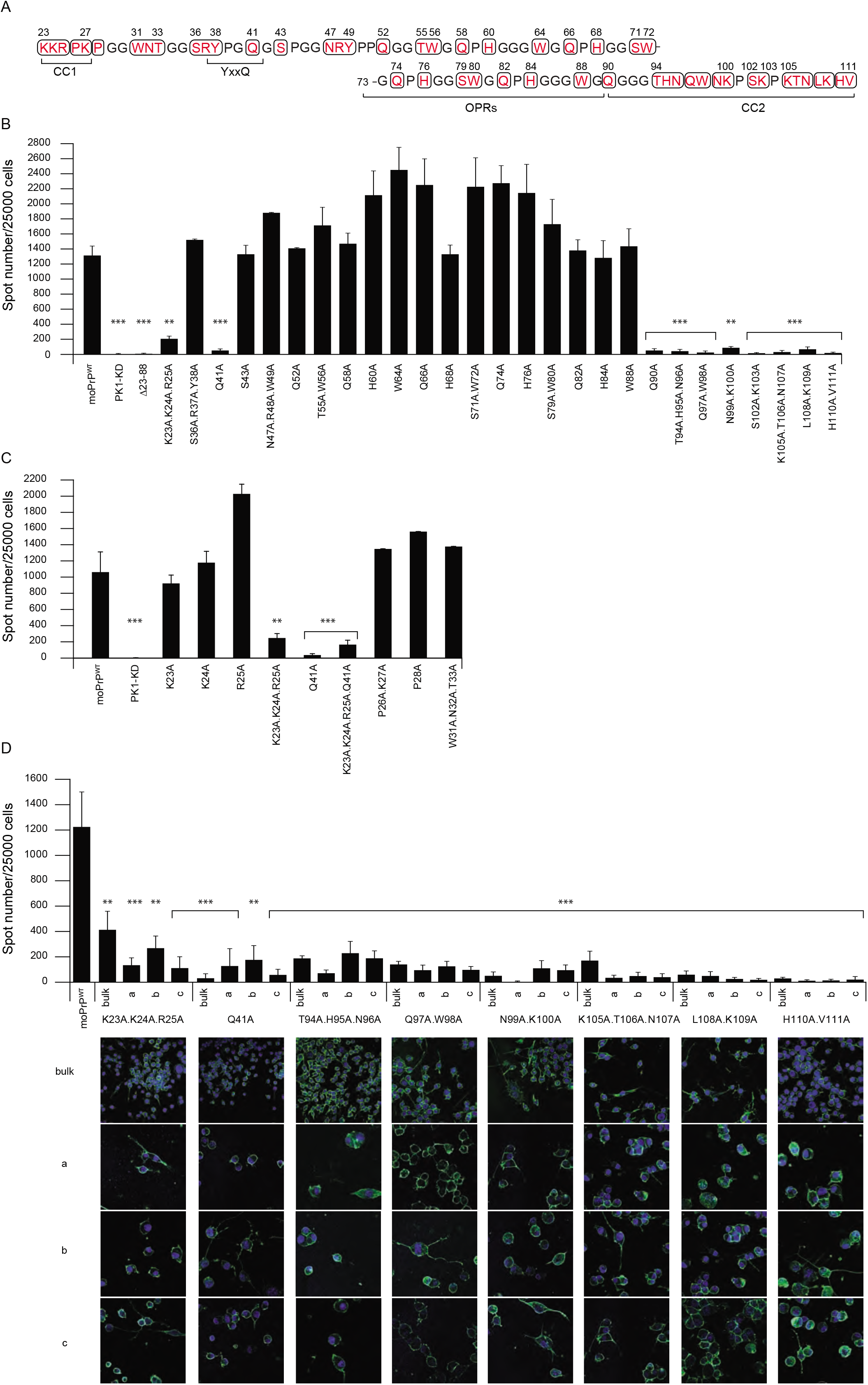
SCA for N-terminal alanine mutations in moPrP. A. All residues other than proline, glycine and existing alanine (black) within the N-terminal 23-111 region were targeted for mutagenesis (red, circled). B. SCA data for split 6, post-RML infection [10^-5^ dilution of 10% infected brain homogenate] of PK1, PK1-KD and bulk cultures of PK1-KD cells reconstituted with the indicated moPrP mutants. C. SCA data for mutants within region 23-41. D. SCA data and immunofluorescence images of single cell clones of PK1-KD cells after reconstitution with the indicated moPrP mutants showing PrP expression. DAPI, nuclear stain; mouse PrP, green. For SCAs, significance was calculated in a one-way ANOVA with Bonferroni correction for multiple comparisons to PK1-KD cells reconstituted with moPrP^WT^. It is indicated by *** for *p*≤0.0001, ** for *p* ≤0.0002, * for *p* ≤0.005.

Rather unexpectedly, no mutation within the OPRs was found to reduce prion propagation (Fig. 3B and Fig. EV2B). Alanine mutations tested encompassed all OPRs and also included point mutations Q52A, Q58A, H60A, W64A, Q66A, H68A, Q74A, H76A, Q82A, H84A and W88A, and double mutations T55A.W56A, S71A.W72A and S79A.W80A. PK1-KD cells reconstituted with S43A, the triple mutant N47A.R48A.W49A and the OPR mutants yielded spot numbers ≥ PK1 and PK1-KDmoPrP^WT^ cells (Fig. 3B and Fig. EV2B).

To investigate whether reduced prion propagation in the KKR (23-25) mutant was due to removal of the three charge residues, or mutation of one key residue, single replacements were made to produce K23A, K24A and R25A. When assayed for prion propagation, none of the cells expressing single replacements displayed reduced spot numbers compared to PK1-KDmoPrP^WT^ (Fig. 3C and Fig. EV2C). To determine if KKR (23-25) was the critical domain that regulated prion propagation, or neighbouring residues also contributed to this phenotype, cells expressing mutations P26A.K27A, P28A and W31A.N32A.T33A were generated and challenged with RML prions. None of them reduced prion propagation relative to PK1-KDmoPrP^WT^, delineating KKR (23-25) as the modulator of prion propagation (Fig. 3C and Fig. EV2C). When mutation of KKR (23-25) was combined with mutation of Q41, the effects on RML propagation were not exacerbated (Fig. 3C and Fig. EV2C). These results indicating that the mutation of KKR or Q41 or a combination thereof (K23A.K24A.R25A.Q41A) was not as inhibitory as Δ23-88, suggested that either all the other residues within the 23-88 region contribute slightly to prion formation, or the Δ23-88 deletion perturbs the native folding of PrP (Ostapchenko *et al*, 2008), thereby reducing its availability as a substrate for prion propagation.

Following analysis of aa23-88, we investigated the adjacent aa89-110 CC2 domain’s effect on prion propagation. Remarkably, all mutations within the CC2 region, demarcated by Q90A and H110A.V111A (Fig. 3A), markedly reduced RML propagation. They produced significantly fewer spots compared to PK1-KDmoPrP^WT^ and PK1 cells. Spot numbers for CC2 mutations were above background (PK1-KD and Δ23-88), comparable to Q41 and less than the KKR (23-25) (Fig. 3B and Fig. EV2B). To determine if increasing the available pool of infectious prions would improve the propagation profile of PK1-KD cells reconstituted with mutations at KKR (23-25), Q41 and within CC2, SCA was conducted with a ten-fold higher dose of RML (10^-4^ dilution of 10% infected brain homogenate). Indeed, K23A.K24A.R25A, Q41A and K23A.K24A.R25A.Q41A resulted in higher spot numbers with higher doses of RML innoculum (Fig. EV3A). However, none of the cells expressing CC2 mutants showed significantly increased prion propagation, although a mild increase in spot number was noted for mutation N99A.K100A (Fig. EV3B). All effects on prion propagation for the reported mutants were verified in three independent clonal cell lines isolated from a repeat reconstitution of PK1-KD cells (Fig. 3D and Fig. EV2D). Together these results indicated that mutations at aa23-25 (CC1), 41 (Q41) and 90-111 (CC2) significantly reduced propagation of RML prions. The reduction was greater for the CC2 mutants than KKR (23-25), or KKRQ (23-25, 41) combined.

### Mutation of KKR (23-25), Q41 and the CC2 domain reduce prion infection

To investigate if the prion propagation defect in PK1-KDmoPrP^Ala^ cells was due to reduced susceptibility of these cells to RML prion infection, or their inability to transmit prions to neighbouring cells, PK1-KD cells reconstituted with various mutants were infected with RML as per the standard SCA protocol(Klohn *et al*., 2003). Cells were harvested at split 6, to allow maximum accumulation of *de novo* prions and dilution of the original inoculum and ribolysed. The level of infectivity present in the ribolysed lysates was determined by applying this lysate to highly susceptible PK1 cells via SCA (Fig. 4A and Fig. EV4A). PK1 cells were thus used as reporters of infectivity, for lysates prepared from PK1-KDmoPrP^WT^ (moPrP^WT^) and PK1-KDpLNCX2 (pLNCX2, empty vector control) used as positive and negative controls respectively. PK1 cells infected with lysate from PK1-KDmoPrP^WT^ and PK1-KDS43A, gave spot numbers above 500 at split 6, suggesting similar levels of infectivity were present in these samples. Maximum spot numbers of 200, 100 and 50 spots, were obtained for PK1 cells infected with lysate from PK1-KDmoPrP^Ala^ cells with KKR (23-25), Q41 and CC2 domain mutations (T94A.H95A.N96A, Q97A.W98A, N99A.K100A, K105A.T106A.N107A and L108A.K109A) respectively indicating decreasing infectivity in these samples (Fig. 4A and Fig. EV4A). All PK1 cells infected with lysates from PK1-KDmoPrP^Ala^ cells with CC2 mutations gave spot numbers equal to or fewer than 50, a background number of spots observed for lysates from PK1-KDpLNCX2. This indicated that the prion propagation defect observed in PK1-KDmoPrP^Ala^ cells reconstituted with KKR (aa23-25), Q41, and the CC2 mutants was due to reduced susceptibility of these cells to RML infection.

**Figure 4.**
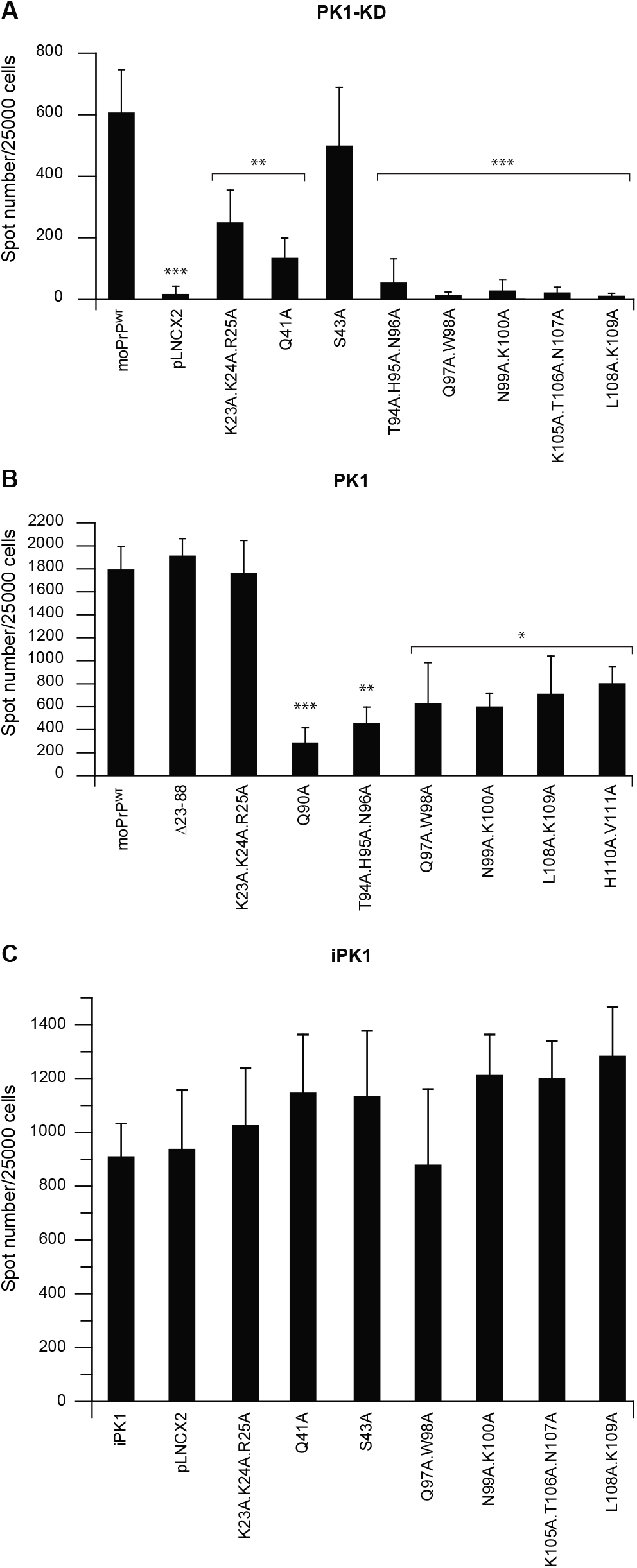
N-terminal residues regulate prion infection. A. PK1-KD cells reconstituted with the indicated mutants were infected with RML (10^-5^ dilution of 10% infected brain homogenate), lysates prepared and applied to susceptible PK1 cells in a SCA. The infectious lysate was used in place of RML to gauge the prion infectivity present in each lysate. B. PK1-cells were stably transduced with retrovirus encoding moPrP with alanine mutations and the resultant cells (expressing both moPrP^WT^ and mutant moPrP) assayed for their sensitivity to RML prions (10^-5^ dilution of 10% infected brain homogenate) in SCA at split 6. C. Chronically prion-infected iPK1 cells were stably transduced with retroviruses encoding moPrP alanine mutants and seeded onto ELISPOT plates for detection of PrP^Sc^ positive spots at split 3 post-transduction. For all SCAs, significance was calculated in a one-way ANOVA with Bonferroni correction for multiple comparisons to PK1-KD cells reconstituted with moPrP^WT^. It is indicated by *** for *p* ≤0.0002, * for *p* ≤0.005 (Yang *et al*, 2022).

### Mutations within the CC2 domain dominantly inhibit prion propagation in PK1 cells

PK1 cells were stably transduced with Δ23-88, KKR (aa23-25) and CC2 mutants and the resulting cells challenged with RML to assess whether these mutations would interfere with prion propagation when co-expressed with wild type PrP^C^. All six mutations tested within CC2 significantly reduced prion propagation, whereas KKR (23-25) and Δ23-88 mutations had no effect (Fig. 4B and Fig. EV4B). PK1 cells expressing CC2 mutants Q97A.W98A, N99A.K100A, L108A.K109A, and H110.V111A yielded slightly higher spot numbers than Q90A and T94A.H95A.N96A at split 6, though all gave considerably fewer spots than PK1Δ23-88 and PK1K23A.K24A.R25A cells, which showed no reduction in prion propagation (Fig. 4B and Fig. EV4B). The lack of prion propagation inhibition by KKR (23-25) and Δ23-88 mutants in PK1 cells suggests that these PrP variants may, as previously mentioned, suffer bioavailability or folding defects (Ostapchenko *et al*, 2008), which prevents them from interfering with prion propagation. In contrast, the CC2 mutants may be avidly recruited to prion fibrils and are thus capable of perturbing their assembly and/or fission. Our finding that Δ23-88 and K23A.K24A.R25A do not inhibit propagation when co-expressed with moPrP^WT^ agrees with previous observations that upon RML infection, moPrP^WT^ can act in *trans* to propagate prions despite the co-expression of a PrP double deletion mutant (Δ23-88 and Δ141–176) in transgenic mice (Supattapone *et al*., 1999) (Supattapone *et al*., 2001). However, mutations within CC2 inhibit prion propagation when co-expressed with moPrP^WT^.

### Mutation of KKR (23-25), Q41 and the CC2 domain inhibit formation of *de novo* prions in uninfected cells but do not inhibit an established prion infection

To determine if the KKR, Q41 and CC2 domain mutations could exert a curing effect in cells with an existing prion infection, these PrP mutants were expressed in chronically prion-infected PK1 (iPK1) cells. iPK1 cells were stably transduced with retroviruses encoding KKR (23-25), Q41 and four CC2 [QW (97-98), NK (99-100), KTN (105-107) and LK (108-109)] mutants and the resulting cells assessed for levels of PrP^Sc^ by ELISPOT. No marked differences were found in the PK-resistant spot numbers between iPK1pLNCX2 (empty vector control) or iPK1 cells transduced with KKR (23-25), Q41 or CC2 mutants, indicating that mutations in these regions exerted no control over an established prion infection as their expression in iPK1 cells did not, ‘cure’ the cells of infection (Fig. 4C).

### Are KKR (23-25), Q41 and the CC2 domain required for propagation of 22L, Me7 and MRC2 prions?

To determine if these three regions were also required for propagation of other prion strains, CAD-2A2D5 (CAD5) cells, that are susceptible to RML, ME7, 22L and the BSE-derived 301C strains of prions (Qi *et al*, 1997) (Mahal *et al*, 2007) were utilised. Endogenous PrP^C^ was silenced using shRNA8, as used for silencing PK1 cells. Bulk cultures of stably transduced cells were negatively sorted for cell surface expression of PrP^C^ by fluorescence-activated cell sorting (Emma Quarterman and Gigi Yang, unpublished work). The sorted cells were single cell cloned; 24 clones were isolated, expanded into cell lines, reconstituted with full length moPrP^WT^ and challenged with 22L and MRC2 (a mouse-adapted BSE strain) (Lloyd *et al*., 2004a) to identify a clone that regains full susceptibility upon reconstitution. This identified clone CAD5-KDB3, that has highly reduced levels of PrP RNA (<1% of the level in CAD5 cells), essentially no detectable cell surface PrP^C^ expression and can propagate 22L, MRC2 and ME7 prions upon reconstitution with moPrP^WT^ (Fig. 5 A&B and 6A, Figs. EV5&6). CAD5-KDB3 cells are referred to as CAD5-KD from hereon as the only subclone of CAD5 cells used for reconstitution experiments in this study.

**Figure 5.**
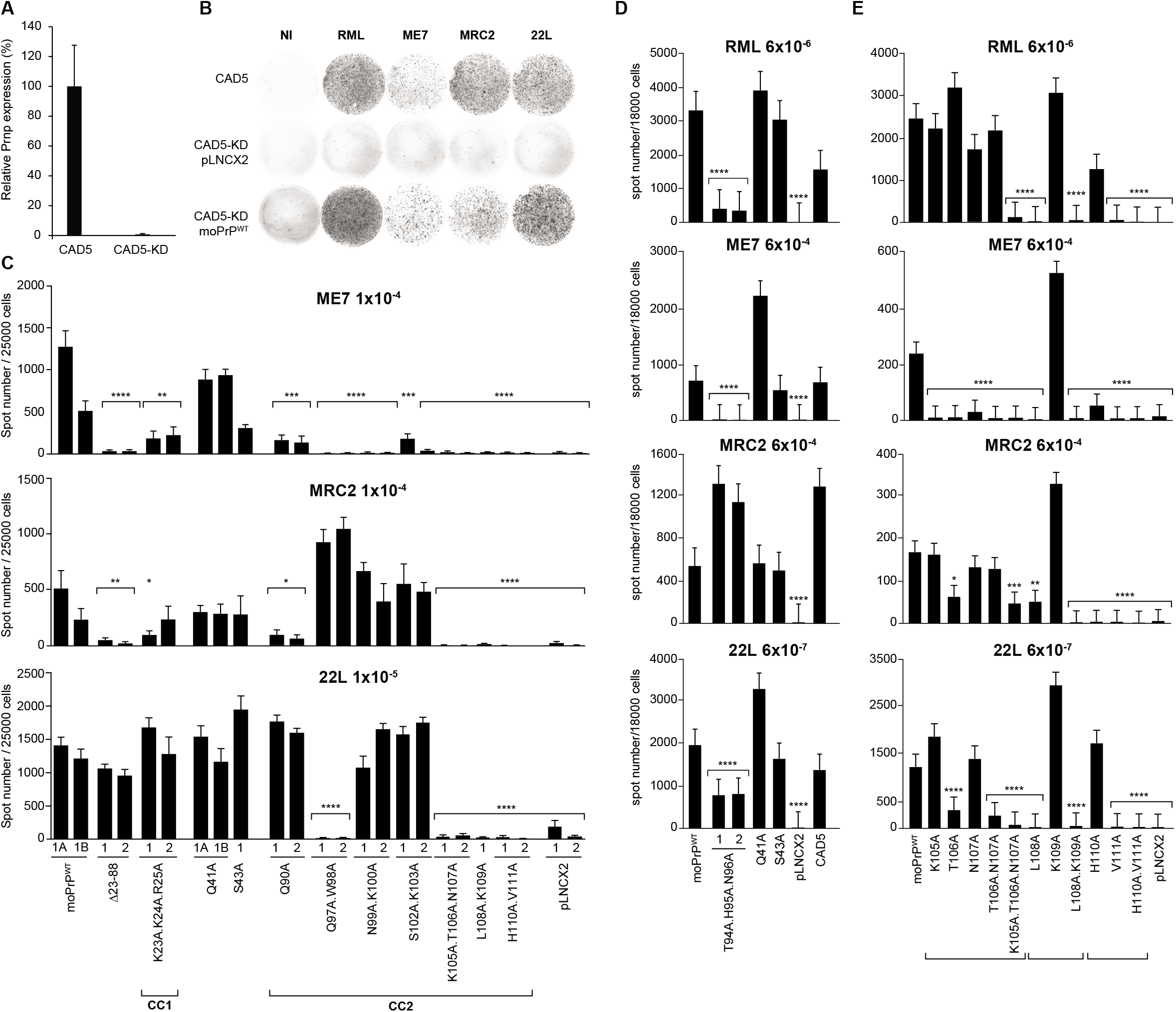
SCA for N-terminal alanine mutations in CAD5-KD cells. A. Expression of *Prnp* in CAD5-KD cells was measured relative to CAD5 cells by RT-PCR. The reactions were carried out in triplicate; the error bars show the standard deviation. *Prnp* expression in CAD5-KD cells was found to be reduced to <1% of the expression in CAD5 cells. B. Representative images of individual ELISPOT wells at split 6 taken from the expt. presented in D. They show CAD5 and CAD5-KD cells reconstituted with pLNCX2 (empty vector) and moPrP^WT^ after no infection (NI) and infection with RML, ME7, MRC2 and 22L prions. C. SCA data for split 6 after infection of CAD5, CAD5-KD and independently isolated bulk cultures of CAD5-KD cells reconstituted with the indicated moPrP mutants following infection with ME7, MRC2 and 22L prions respectively. Cultures A and B are the same bulk culture but at different serial passages. Data are shown as mean ± SD, ****P < 0.0001, ***P < 0.001, **P < 0.01, *P < 0.05. Statistical analyses were performed using an unpaired 2-tailed t-test in GraphPad Prism 7.0 to CAD5-KD cells reconstituted with moPrP^WT^ bulk culture 1B. D and E. SCA data for split 6 after infection of CAD5, CAD5-KD and bulk cultures of reconstituted CAD5-KD cells following infection with RML, ME7, MRC and 22L prions. Data are shown as mean ± SD, ****P < 0.0001, ***P < 0.001, **P < 0.01, *P < 0.05. Statistical analyses were performed using an unpaired 2-tailed t-test in GraphPad Prism 7.0 to CAD5-KD cells reconstituted with moPrP^WT^.

**Figure 6.**
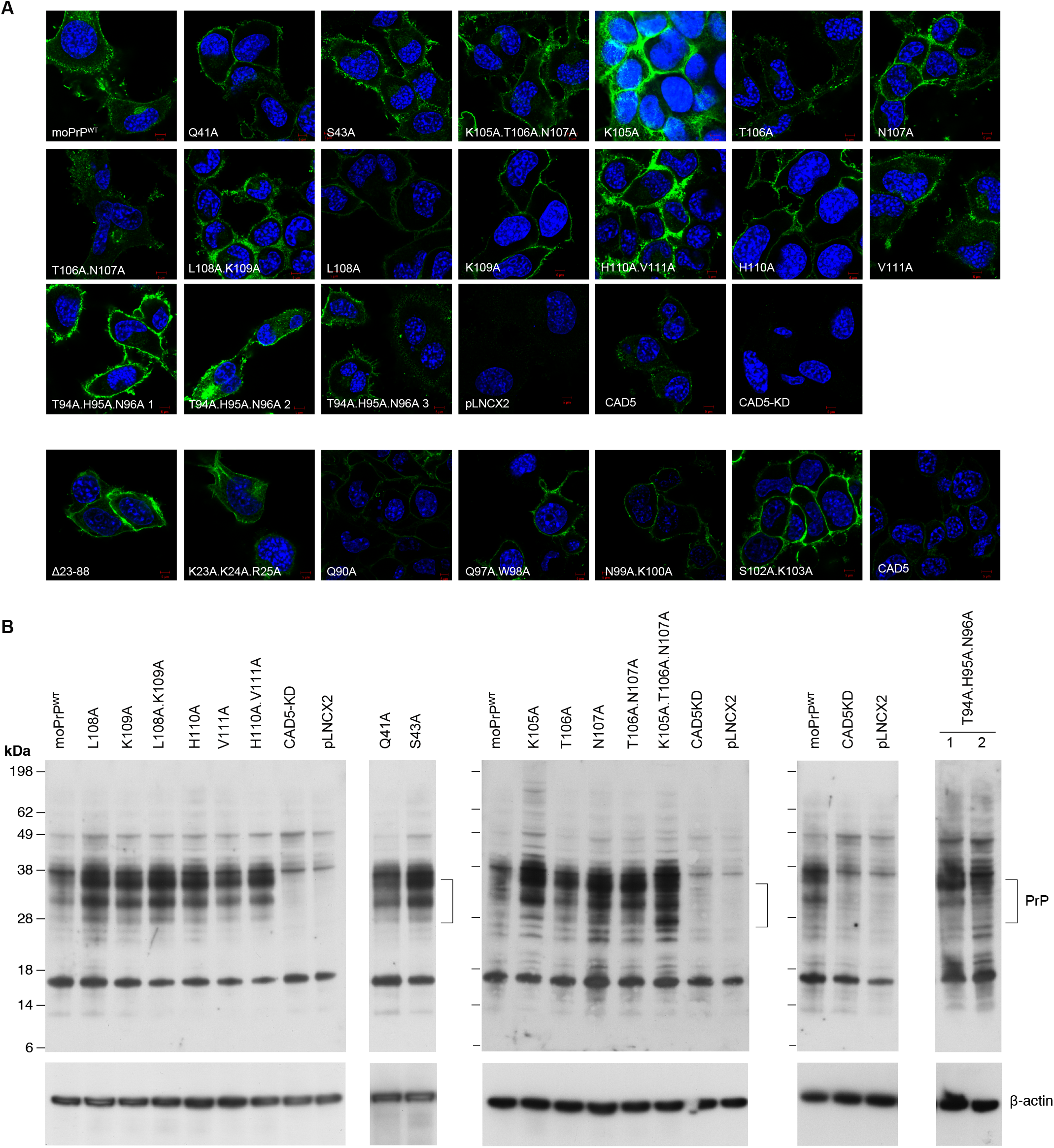
Analysis of PrP expression in reconstituted CAD5-KD cells. A. Laser-scanning microscopy images of bulk cultures of CAD5, CAD5-KD and CAD5-KD cells after reconstitution with the indicated moPrP mutants. The analysis was carried out in two batches as indicated. DAPI, nuclear stain; mouse PrP, green. B. Western blot analysis of PrP expression in bulk cultures of CAD5, CAD5-KD and CAD5-KD cells reconstituted with the indicated mutants.

CAD5-KD cells were stably reconstituted with Δ23-88, KKR, Q41 and CC2 mutants and drug resistant colonies pooled to prepare bulk cultures. Fig. 5C&D and Fig. EV6&7 show representative experiments where independently derived bulk cultures for each mutant were challenged in turn with ME7, MRC2 and 22L prions simultaneously. The results for the ME7 strain (Fig. 5C&D) were akin to those with RML albeit with two differences. ME7 propagation was unaffected by mutation of Q41, whereas mutation of S43 reduced spot numbers, which increased at splits 5 and 6, indicating a lower level of propagation. In contrast, the results for the 22L strain (Fig. 5C&D) showed that mutation of KKR (23-25) or Q41 did not affect propagation, as spot numbers were similar to moPrP^WT^. Cells reconstituted with Δ23-88 were slightly reduced in propagation of 22L in comparison to moPrP^WT^, but exhibited an increase in spot number at three consecutive splits, indicating propagation of 22L prions. Results for CC2 domain mutants were more surprising. Mutants Q90A, N99A.K100A and S102A.K103A supported propagation of 22L whereas mutants Q97A.W98A, K105A.T106A.N107A, L108A.K109A and H110A.V111A did not. Mutant T94A.H95A.N96A supported a reduced level of propagation of 22L (Fig. 5D). The MRC2 strain (Fig. 5C&D) generally produced lower spot numbers than the other three strains. Mutation of Q41 and S43 did not affect propagation whereas Δ23-88 clearly inhibited propagation. The results for CC2 mutants were similar to those with 22L. However, in this case mutants Q97A.W98A, N99A.K100A and S102A.K103A supported MRC2 propagation whereas mutants Q90A, K105A.T106A.N107A, L108A.K109A and H110A.V111A were unable to support propagation. Cells transduced with T94A.H95A.N96A propagated MRC2 even more efficiently than cells reconstituted with full-length mouse PrP (moPrP^WT^). Rather surprisingly, CAD5-KD cells reconstituted with mutant Q41 were found to support propagation of RML, whereas this mutant did not support RML propagation in PK1-KD cells (compare Fig. 5D with Fig. 3B&C). Thus, glutamine 41 is required for propagation of RML in PK1 cells but not in CAD5 cells.

To identify which amino acids within the 105-111 region were critical for propagation of the four-prion strains, each amino acid was individually mutated to alanine. Mutant T106A.N107A was also prepared since aa105-107 (KTN) had originally been mutated together. All of the mutants were used to stably reconstitute CAD5-KD cells (Fig. 5E) and derive bulk cultures of drug resistant colonies that were challenged in turn with RML, ME7, 22L and MRC2 prions simultaneously in SCA; a representative experiment is shown in Fig. 5E and Fig. EV7. The results showed that mutation of K109 did not affect propagation of all four prion strains, whereas mutation of K105 did not affect propagation of RML, MRC2 and 22L; mutation of H110 fully supported propagation of 22L but RML only at a reduced level. In contrast, mutation of L108 and V111 severely impacted propagation of all four prion strains. Mutation of L108 and V111 alone was as effective as mutation of LK (108-109) or HV (110-111) except L108A.K109A was more inhibitory for MRC2 than L108A. Only when K105.T106.N107 were mutated to alanine together, did they impede propagation of all four prion strains.

To confirm that the observed effects on prion propagation were a direct consequence of the mutations within PrP, protein expression in CAD5-KD cells expressing these mutants was examined by western blotting and laser-scanning confocal microscopy. The results in Fig. 6 A&B show that although expression was variable, all the mutants were generally expressed at levels higher than endogenous PrP^C^ and predominantly on the cell surface. Moreover, the mutants exhibited the same pattern of three bands corresponding to endogenous PrP^C^ that is non-, mono- and di-glycosylated in the parental CAD5 cells. This was consistent with previous studies documenting that although PrP^C^ is necessary for replicating infectivity (McNally *et al*, 2009), the actual level of expression is not important. Enari *et al* found that N2a/Bos2 cells, a clone of prion-susceptible cells, expressed PrP^C^ at the same low level as the parental N2a cells, whereas a resistant cell line expressed it at a 10 times higher level (Enari *et al*, 2001). Moreover, over-expression of PrP did not increase susceptibility either, indicating that it is a prerequisite for prion propagation, but other factors are also essential (Enari *et al*., 2001).

Taken together, the SCA data showed that amino acids 105.KTN.107, L108 and V111 within CC2 domain were required for propagation of all four prion strains tested: RML, ME7, 22L and MRC2. Other N-terminal residues either had no effect or only affected propagation of select strains. We have also identified a mutant that behaves differently depending on the cell context. Q41A severely impacted RML propagation in reconstituted PK1-KD cells but had no effect in CAD5-KD cells. Overall, the profile of N-terminal residues required for propagation of ME7 and RML were similar, whereas those required for 22L were very different from these two strains and MRC2 shared features with all three.

## Discussion

This study combined the development of mouse PK1-KD and CAD5-KD cell lines in which endogenous PrP has been stably silenced, such that they are refractory to prion infection while regaining full susceptibility upon reconstitution with mouse PrP^C^, with a systematic alanine replacement mutagenesis of the N-terminal 23-111 region of mouse *Prnp* to identify residues that are required for highly efficient prion propagation. We have shown that mutation of leucine 108 and valine 111 alone or simultaneous mutation of lysine 105, threonine 106 and asparagine 107, severely impacts propagation of RML, ME7, 22L and MRC2 mouse prion strains. Mutation of other N-terminal residues including the octapeptide repeats and CC1 domain either had no effect or only affected propagation of select prion strains. Replacements in the CC2 domain including aa105-111 dominantly inhibited prion propagation in the presence of endogenous PrP^C^ whilst other changes were not inhibitory. None of the mutants including aa105-111 blocked prion propagation when expressed in chronically prion-infected cells. Together, these results indicate that efficient prion propagation is dependent upon leucine 108 and valine 111 individually or lysine 105, threonine 106 and asparagine 107 simultaneously, acting at the infection stage.

N-terminal residues 23-31 comprising the polybasic CC1 domain of the prion protein have been implicated in endocytosis via clathrin-coated pits (Shyng *et al*, 1995) (Ostapchenko *et al*., 2008) (Fehlinger *et al*, 2017), neurotoxicity (Hara & Sakaguchi, 2020) (Turnbaugh *et al*, 2011), prion conversion (Turnbaugh *et al*., 2012) (Hara & Sakaguchi, 2020) as well as association with amyloid-β oligomers (Chen, Yadav et al. 2010; Fluharty, Biasini et al. 2013). *Prnp^0/0^* mice reconstituted with Δ23-31 show prolonged survival after inoculation with RML and accumulate lower levels of PK-resistant PrP (Turnbaugh *et al*., 2012). Our results showed that for RML propagation, the critical residues within the CC1 domain were K23.K24.R25 and simultaneous replacement of all three residues was required to reduce the susceptibility to RML infection (Fig. 4A), but this was reversed by co-expression of wild-type PrP^C^ (Fig. 4B).

Mutation of Q41 exhibited reduced RML propagation in PK1 cells, whereas none of the neighbouring residue replacements, namely SRY (36-38) and S43 had an effect (Fig. 3B). There is no previous evidence implicating Q41 as a site for modulating prion propagation, or indeed for any other prion function. Of the alanine-mutants generated within aa23-88, 14 were within the OPR, each of which propagated RML at levels ≥ moPrP^WT^ (Fig. 3B) suggesting that the OPR domain does not play a critical role in regulating RML propagation. Another possible interpretation of our finding that no single OPR mutation affected propagation may be that our alanine mutagenesis strategy replaced histidines individually rather than replacing all simultaneously. Previous studies have suggested that the OPRs play only a limited role in disease pathogenesis (Yamaguchi *et al*., 2012) but supernumerary insertions of between one and nine additional OPRs, increase the risk of developing disease, with most cases showing an earlier onset (Stevens *et al*., 2009). Deletion of residues 23-88 abrogated RML propagation (Fig. 2C) yet only mutations at KKR (23-25) and Q41 within this region were found to be inhibitory (Fig. 3B). When these two mutations were combined to determine any synergistic effects, none were observed; the results were an average of the individual mutations rather than additive, indicating that KKR (23-25) and Q41 modulate the same pathway probably at an early stage of infection, rather than at a later stage once infection has been established (Fig. 4C).

Minimal alanine substitutions (up to three residues) in the CC2 (90-111) region produced mutants that were well expressed in both PK1-KD and CAD5-KD cells (Fig. 2B, 3D and 6). Q90A defined the most N-terminal position of the protein that showed a tenfold reduction in RML propagation; whereas adjacent mutation W88A did not reduce propagation relative to moPrP^WT^ expressing cells (Fig. 3B). Mutations at the C-terminal end of CC2 (H110A.V111A) also completely eliminated prion propagation. The CC2 domain mutants also reduced RML propagation when expressed in the presence of endogenous PrP^C^ in PK1 cells, a characteristic of competitive inhibition, whereas KKR (23-25) and Δ23-88 mutations were not inhibitory (Fig. 4B).

The next phase of this study was to examine if the requirement for KKR (23-25), Q41 and CC2 was RML-specific or a general requirement for prion infection and propagation. Since CAD5 cells are susceptible to RML, ME7, 22L and the BSE-derived 301C strain (Qi *et al*., 1997), we reconstituted CAD5-KD cells with each of the mutants and challenged them with 22L, ME7 and MRC2 prions. MRC2 is a mouse-adapted BSE prion strain characterised by a long incubation time and di-glycosylated-dominant PrP^Sc^ (Lloyd *et al*., 2004a) and may represent the same strain as 301C. The results showed that Δ23-88 inhibited propagation of ME7 and MRC2 but 22L was still able to propagate albeit at a slightly lower efficiency, as in Uchiyama *et al*, who found that mouse PrP Δ23-88 restored susceptibility to 22L but not RML prions in *Prnp^0/0^* mice (Uchiyama *et al*, 2014). Similarly, aa23-25 (KKR) were required for propagation of ME7 and probably MRC2 prions but did not affect propagation of 22L prions. Khalifé *et al* have found that deletion of aa23-26 (KKRP) in transgenic mice overexpressing ovine PrP resulted in variable susceptibility to prion infection, depending on the prion strain (Khalife *et al*, 2016), further highlighting aa23-25 as a strain-dependent modulator of prion propagation.

Like for the RML experiments in PK1 cells, all CC2 domain mutants affected propagation of ME7 prions. Mutations within aa94-98 and 105-111 inhibited 22L propagation whereas mutation of Q90, N99.K100 and S102.K103 had no effect. Propagation of MRC2 was inhibited by mutation of Q90 and aa105-111 whereas mutations within aa94-103 had no effect. Taken together these findings indicated that aa105-111 were required for efficient propagation, independent of prion strain (RML, 22L, ME7 and MRC2). The four prion strains can be distinguished from each other by the N-terminal amino acids that are required for their efficient propagation; RML and ME7 are the most similar whereas 22L exhibits the greatest differences to RML and ME7; MRC2 shares features with the other three. Analysis of single alanine replacements within aa105-111 showed that L108A and V111A alone were as effective as L108A.K109A and H110A.V111A for inhibiting prion propagation except L108A.K109A was more inhibitory for MRC2 than L108A. Neither L108 nor V111 are conserved between mice, hamsters and humans and are part of the epitope for the 3F4 antibody that can distinguish hamster and human PrP^C^ from mouse PrP^C^. PK1-KD cells reconstituted with moPrP tagged with the 3F4 epitope do not support propagation of RML prions (Fig. EV8).

The recent structures of mouse brain-derived RML and ME7 prion fibrils (Manka *et al*., 2022a) (Manka *et al*., 2022b) show that a large portion (aa94-111) of the CC2 domain, which is disordered in PrP^C^, becomes ordered in these prion fibrils (together with the LCR region), forming the first two β-strands in β-sheet-rich assemblies from both strains (Fig. 7). PrP residues that we identified as key for propagation of all four mouse prion strains (105.KTN.107, L108 and V111), are all located around the second β-strand. The two key hydrophobic residues (L108 and V111) and the aliphatic portion of T106 are part of large hydrophobic cluster 1 (T106, L108, V111, I138, F140) that stabilises the PrP fold in each fibril (Fig. 7). K105 contributes to the major basic patch on the surface of each fibril, which is thought to be an interaction hub for disease-associated PrP, as identified using motif grafted antibodies (Solforosi *et al*, 2007) (Abalos *et al*., 2008). L108 is also one of the two amino acids (L108 and T189) that define the *Prnp^a^* allele associated with a short incubation time upon prion infection in a range of inbred mouse strains (Lloyd *et al*, 2004b). Taken together, our data suggest that amino acid side chains that favour longitudinal cross-β stacking, coupled with strong lateral hydrophobic interaction via side chains of hydrophobic cluster 1 (Fig. 7), are required for the specific ordering of the CC2 region.

**Figure 7.**
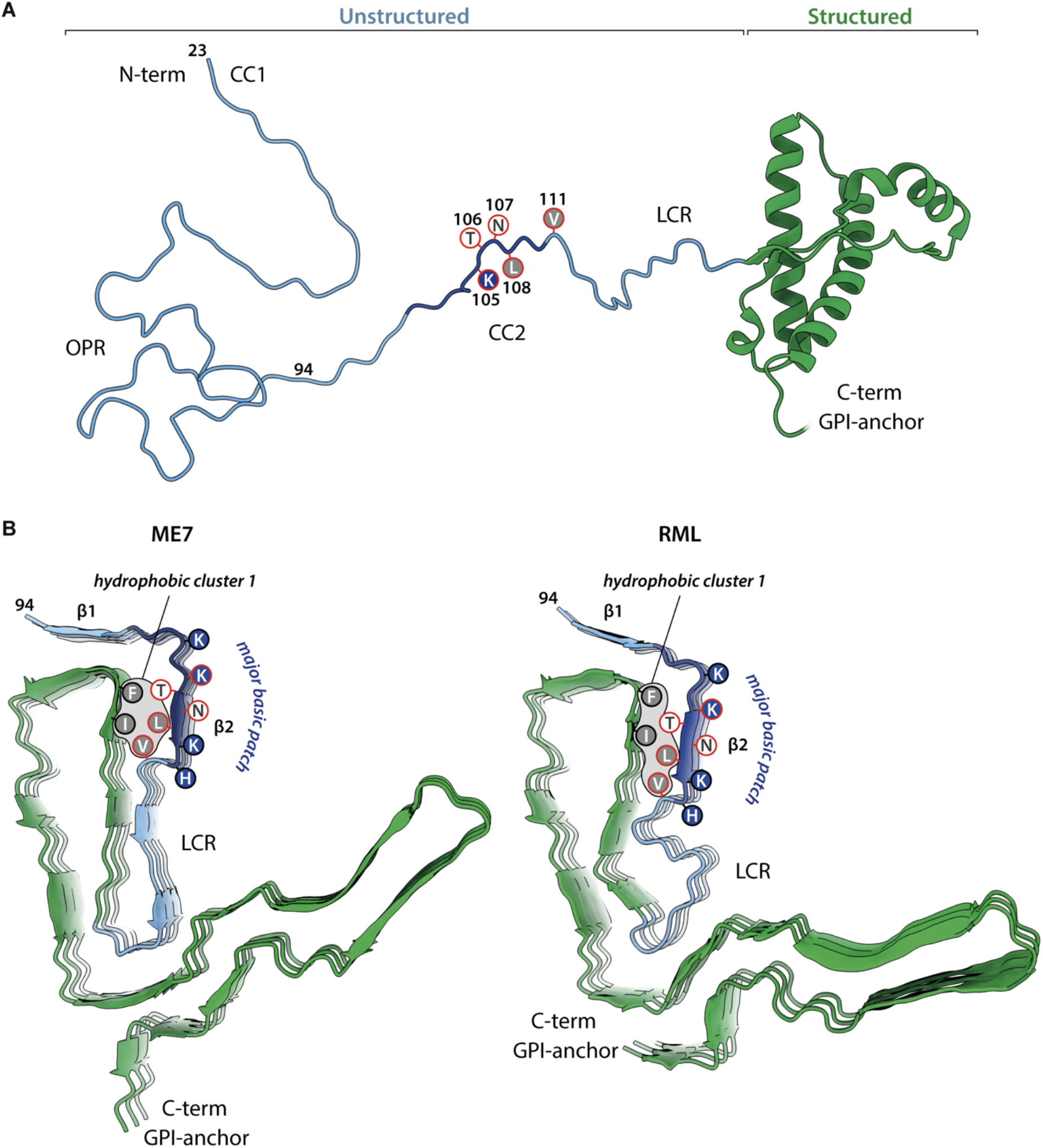
Mapping residues critical for prion propagation on PrP and prion fibril structures. A. Ribbon model of mature mouse PrP^C^ (residues 23-230, excluding post-translational modifications) built in UCSF Chimera (Pettersen *et al*, 2004) using an X-ray structure of moPrP^C^ (pdb ID: 4H88) (Sonati *et al*, 2013). Positions of amino acid side chains found to be critical for prion propagation (basic, navy blue; neutral, white; hydrophobic, grey) are indicated. CC1, Charge Cluster 1; OPR, Octapeptide Repeats; CC2, Charge Cluster 2; LCR, Low-Complexity Region; GPI, glycosylphosphatidylinositol. B. Mouse prion fibril structures from RML (pdb ID: 7QIG) (Manka *et al* 2022) and Me7 (pdb ID: 8A00) (Manka *et al* 2022) strains (3 subunits, ribbon representation, coloured as in (A)). Amino acid side chains found to be critical for prion propagation (marked with red circles) in the context of surrounding residues, coloured as in (A) are indicated. Major internal hydrophobic cluster 1 that contributes to PrP fold stability is indicated together with the major basic patch (navy blue). β-strands 1 and 2 are labelled.

Distinct prion strains present various degrees of sensitivity to mutations in this critical CC2 region. For example, single K105A or T106A or N107A mutations are sufficient to abrogate propagation of ME7, while having no effect on the propagation of RML (Fig. 5E). The atomic structures of ME7 and RML fibrils reveal structural differences between these strains, which include distinct interactions between T106 and neighbouring residues of hydrophobic cluster 1 (Fig. 7B). Only three simultaneous alanine replacements in the 105.KTN.107 segment abrogate propagation of all mouse prion strains tested. The structures of 22L and MRC2 assemblies are not yet known. Our data suggests that their ordering of the CC2 region will be similar overall, but with subtle differences that translate to the differential sensitivity to alanine replacements. Future experiments will be directed at determining if 105.KTN.107 is sufficient for binding disease-associated PrP to initiate seeded protein polymerization or if another (co-)factor is required for prion propagation.

## Materials and Methods

### Construction of plasmid DNAs

pBluescript SK+ plasmid vector containing the full length ORF for mouse PrP was used as the template DNA. All residues within aa23-110 except glycine and proline were mutated to alanine in blocks of one, two or three amino acids, using the Stratagene QuikChange® site-directed mutagenesis system (Agilent Technologies, Santa Clara CA9051, USA). Mutations to alanine were designed using the moPrP protein sequence (UniProtKB entry P04925 NCBI Reference Sequence NM_011170.3). Codon GCC was selected for alanine replacement based on codon bias in the mouse genome. Positions at which native residues were proline or glycine were not targeted for mutagenesis. Sequence-verified mutated ORFs were inserted into the pLNCX2 retroviral vector (Clontech Takara, Mountain View CA 94043, USA). The siRNA target sequences were used to generate 64mer DNA oligonucleotides in accordance with the Oligoengine template design (www.oligoengine.com). The oligonucleotide pairs were annealed and inserted into the pRetroSuper silencing vector.

### Retroviral expression

pRetrosuper shRNA and pLNCX2 constructs were packaged as ecotropic retroviruses in Phoenix ecotropic cells (ATCC, LGC Standards, Middlesex, UK) and used to stably transduce PK1 and CAD5 cells and derivatives thereof. The efficiency of stable transduction of CAD5-KD cells was increased by pseudotyping the ecotropic retroviruses using the vesicular stomatitis virus G (VSV-G) protein. Stable transduction of cells with pLNCX2 retroviruses was selected using 300µg/ml G418 for PK1 and 400µg/ml for CAD5 cells respectively. Stable silencing of endogenous PrP was achieved through stable expression of pRetrosuper shRNA constructs by selecting for puromycin resistance at 4µg/ml for PK1 and 2µg/ml for CAD5 cells respectively.

### Scrapie Cell Assay

SCA was carried out as previously described (Klohn *et al*., 2003). Cells were plated at 18,000 cells/well of a 96-well plate, infected with prion infected brain homogenates the following day and grown for three weeks with 1:8 biweekly splits. At splits 4, 5 and 6, cell suspensions equivalent to 25,000 PK1 or 18,000 CAD5 cells were plated onto activated ELISPOT plates and probed for PK-resistant PrP using anti-PrP ICSM18 antibody (D-Gen Ltd, UK). Cells positive for PK-resistant PrP were quantified initially using WellScan software (Imaging Associates, Oxfordshire, UK) and more recently a Bioreader 7000F Alpha (BIOSYS GmbH, Germany).

### Western blotting

PK1 cell lysates were prepared from frozen pellets by resuspending in 150mM NaCl, 50mM TrisHCl pH 7.5, 0.5% Triton-X-100, 0.5% sodium deoxycholate, 1mM EDTA and 40U/ml benzonase. 25μg of protein was fractionated on 16% Novex® Tris-Glycine mini gels, transferred to PVDF membrane and probed using anti-PrP ICSM18 antibody (D-Gen LTD, London, UK).

CAD5 cell lysates were prepared from frozen cell pellets enriching for membrane bound proteins. Pellets were resuspended in 10mM phosphate buffer (P5244, SigmaAldrich), incubated on ice at 4°C for 5 mins followed by centrifugation at 15,000g for 15 mins. The pellet comprising the membrane fraction was resuspended in D-PBS and treated with benzonase (1-2μl, 25KU equivalent to >250units/μl) at room temperature for 15 mins. An equal volume of 2x sample buffer (125mM TrisHCl pH6.8, 20 % v/v glycerol, 4 % w/v SDS, 4 % v/v 2-mercaptoethanol, 8mM 4-(2-aminoethyl)-benzene sulfonyl fluoride and 0.02 % w/v bromophenol blue) was added and the samples boiled for 10 mins, followed by centrifugation at 15000g for 1 min. Supernatant was removed and protein concentration determined by the Bradford Assay.

15-25μg of protein was fractionated on 12% BisTris NUPAGE gel (NP0341, ThermoScientific), transferred to PVDF and probed overnight at room temperature using anti-PrP ICSM35 antibody (0.2μg/ml, D-Gen LTD, London, UK). Antigen-antibody complexes were identified using goat anti-mouse AP (1:10,000 dilution of A2179, Sigma-Aldrich) and CDP-*Star*™ Substrate (T2146, ThermoScientific).

### Immunofluorescence analysis

20,000 PK1 cells and derivatives thereof were seeded on sterile poly-L-lysine coated coverslips, grown at 37°C and fixed using 4% w/v paraformaldehyde. PrP expression was determined using the anti-PrP ICSM18 antibody and visualised using AlexaFluor 488-conjugated goat anti-mouse secondary antibody.

50,000 CAD5 cells or derivatives thereof were seeded in each well of a chamber slide and grown at 37°C for 3 days before fixation in 3.7% w/v paraformaldehyde. Anti-PrP ICSM18 (1μg/ml) diluted in D-PBS containing 25% w/v superblock and 10% v/v penicillin streptomycin was added to each well of the chamber slide and incubated overnight at 4°C. After extensive washing, the chamber slides were incubated again overnight at 4°C with AlexaFluor 488-conjugated anti-mouse IgG (H+L) (115-545-116, Stratech Scientific LTD) in D-PBS containing 25% w/v superblock and 10% v/v penicillin streptomycin and DAPI (4′, 6-diamidino-2-phenylindole). After thorough washing, the chamber slides were imaged using a Zeiss LSM710 laser-scanning microscope, equipped with a 63x objective (Carl Zeiss, Cambridge, UK).

### Cell viability assay and dot blot for moPrP expression

Cells were seeded at 2.5 x 10^4^ cells/well in flat-bottomed 96-well plates and lysed with CellTiter-Glo® reagent according to the manufacturer’s instructions (Promega Corporation, Madison, WI 53711, USA). Plates were set to shake for 2 mins and incubated for 10 mins at room temperature prior to recording luminescence (integration time = 100 ms). Lysed cells were transferred to activated nitrocellulose using a dot blot manifold, at two cell densities. PrP levels were determined using anti-PrP ICSM18 primary and goat anti-mouse IRDye® 800CW infrared dye secondary antibody and quantified using an Odyssey infrared scanner.

### RT-PCR analysis of *Prnp* gene expression in CAD5 cells

10,000 cells were lysed using the Taqman® Gene Expression Cells to CT™ kit (Ambion, Life Technologies) and the RNA reverse transcribed according to manufacturer’s protocol. The resulting cDNA was assayed using Fam-labelled Prnp Taqman® assay (Mm00448389_m1, ThermoFisher) duplexed with VIC-labelled GAPDH endogenous control (Mm00712869_m1, ThermoFisher). Reactions were carried out in triplicate using an Applied Biosystems 7500 Fast Real-Time PCR machine with cycling conditions: 94°C 15mins; 95°C 15s, 60°C 60s for 40 cycles.

## Supporting information

Extended View Figs. 1-8

## Acknowledgements

We are indebted to Emma Quarterman and Gigi Yang for providing cultures of CAD5 silenced cells that had been negatively sorted for cell surface PrP^C^ expression, Mitali Patel for undertaking the experiment presented in Fig. EV8 and Jonathan Wadsworth for helping us to develop the hypotonic lysis protocol for extracting membrane-associated proteins. We thank Ray Young and Richard Newton for graphics. Work was funded by the UKRI Medical Research Council.

## Author contributions

SB, AC, JC and PSJ conceived and designed the research; SB, PA, CS, CB, MLDR and PSJ performed the research; SB, PA, AC, JC and PSJ analysed the data; SM provided insights into the prion fibril structures; SB, SM, PK, JC and PSJ wrote the paper.

Figure EV1. **SCA output**

A. Double log plot of spot number against tissue culture infectious units (TCIU).

B. Serial dilution of RML homogenate and calculated SCEPA data for TCIU.

C. SCA data from PK1 cells, PK1-KD cells, and PK1-KD cells reconstituted with pLNCX2, moPrP^WT^ and Δ23-88 PrP.

D. Schematic of RML serial dilution in 96-well plate format for six concentrations shown in B.

E. ELISPOT revelation of a positive well.

F. ELISPOT revelation of a negative well.

Figure EV2. **Complete SCA data for PrP mutants**

Complete SCA data showing spot number for splits 4, 5 and 6 post prion-infection.

For clarity however, significance is only shown for data at split 6. *** denotes *p*≤0.0001, ** for *p* ≤0.0002, * for *p* ≤0.005. Significance was calculated in a one-way ANOVA with Bonferroni correction for multiple comparisons to PK1-KD cells reconstituted with moPrP^WT^.

A. SCA data showing spot numbers for PK1, PK1-KD and reconstituted PK1-KD cells.

B. SCA data for PK1, PK1-KD and bulk cultures of PK1-KD cells reconstituted with the indicated moPrP mutants.

C. SCA data for mutants within region 23-25 and Q41.

D. SCA data for bulk and single cell clones of PK1-KD cells after reconstitution with the indicated moPrP mutants.

Figure EV3. **Increased dose of infectious innocula does not rescue propagation of CC2 alanine mutants.**

A. PK1 and PK1-KD cells expressing moPrP with alanine mutations in region CC1 and Q41 report higher spot numbers per 25,000 cells with a ten-fold increase in infectious innocula [10^-4^ dilution of 10% infected brain homogenate].

B. A similar dose response to RML innocula was not observed for alanine mutations in CC2 region.

Significance was calculated in a one-way ANOVA with Bonferroni correction for multiple comparisons to PK1-KD cells reconstituted with moPrP^WT^. It is only shown for data at split 6. *** denotes *p*≤0.0001, ** for *p* ≤0.0002, * for *p* ≤0.005.

Figure EV4. **Complete SCA data for N-terminal residues regulate prion infection**

A. SCA data at splits 4, 5 and 6 post-infection for results in Fig. 4A.

B. SCA data at splits 4, 5 and 6 post-infection for results in Fig. 4B.

Significance was calculated in a one-way ANOVA with Bonferroni correction for multiple comparisons to PK1-KD cells reconstituted with moPrP^WT^. It is only shown for data at split 6. *** denotes *p*≤0.0001, ** for *p* ≤0.0002, * for *p* ≤0.005.

Figure EV5. **Serial dilution of 22L, ME7 and MRC2 on CAD5 and CAD5-KD cells**

SCA data at splits 4, 5 and 6 post-infection.

Figure EV6. **Complete SCA data for mutants after infection with 22L, MRC2 and ME7 prions**

SCA data for splits 4, 5 and 6 post-infection of CAD5, CAD5-KD and bulk cultures of CAD5-KD cells reconstituted with the indicated moPrP mutants presented in Fig. 5C.

Figure EV7. **Complete SCA data for mutants after infection with RML, ME7, MRC2 and 22L prions**

SCA data for splits 4, 5 and 6 post-infection of CAD5, CAD5-KD and bulk cultures of CAD5-KD cells reconstituted with the indicated moPrP mutants presented in Fig. 5 D and E.

Figure EV8. **Mouse PrP^3F4^ does not support RML propagation**

PK1-KD cells were stably transduced by retroviral infection with moPrP^WT^ or moPrP^3F4^. Bulk cell cultures were challenged with 3.3 x 10^-4^ dilution of RML infected brain homogenates in an automated SCA. Cells reconstituted with moPrP^3F4^ were found to stably express 3F4 tagged mouse PrP but were unable to support propagation of RML whereas moPrP^WT^ expressing cells propagated RML.

