## Extended View Figs. 1-8 for "Prion propagation is dependent on key amino acids in Charge cluster 2 within the prion protein"

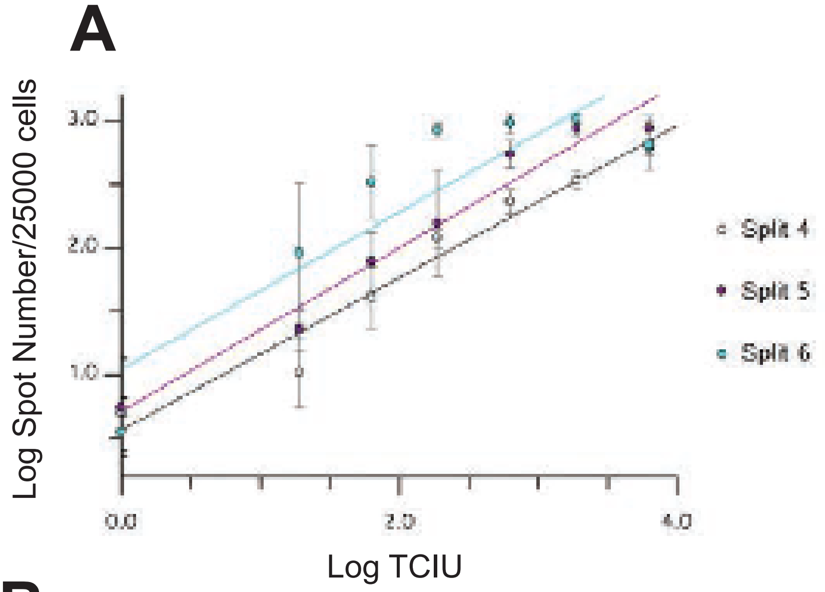


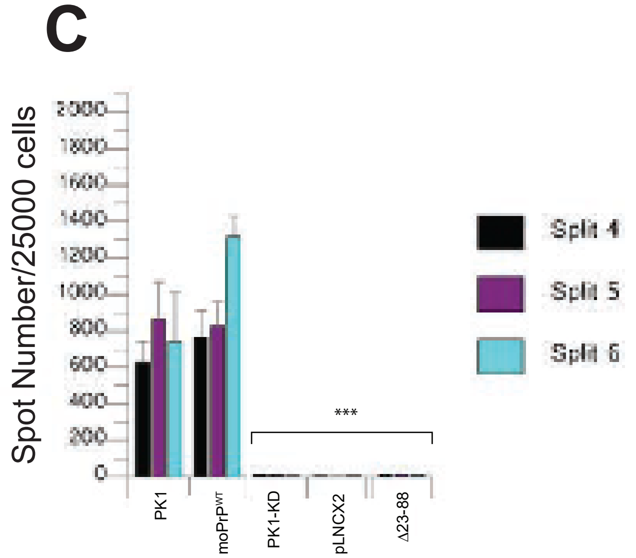


**B**

| Dilution | $3x10^{-5}$ | $1x10^{-5}$ | $3x10^{-6}$ | $1x10^{-6}$ | $3x10^{-7}$ | $1x10^{-7}$ |
|---|---|---|---|---|---|---|
| TCIU | 6309 | 1893 | 631 | 189 | 63.1 | 18.9 |

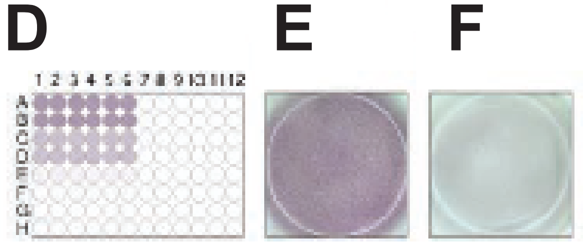


Extended Fig 1

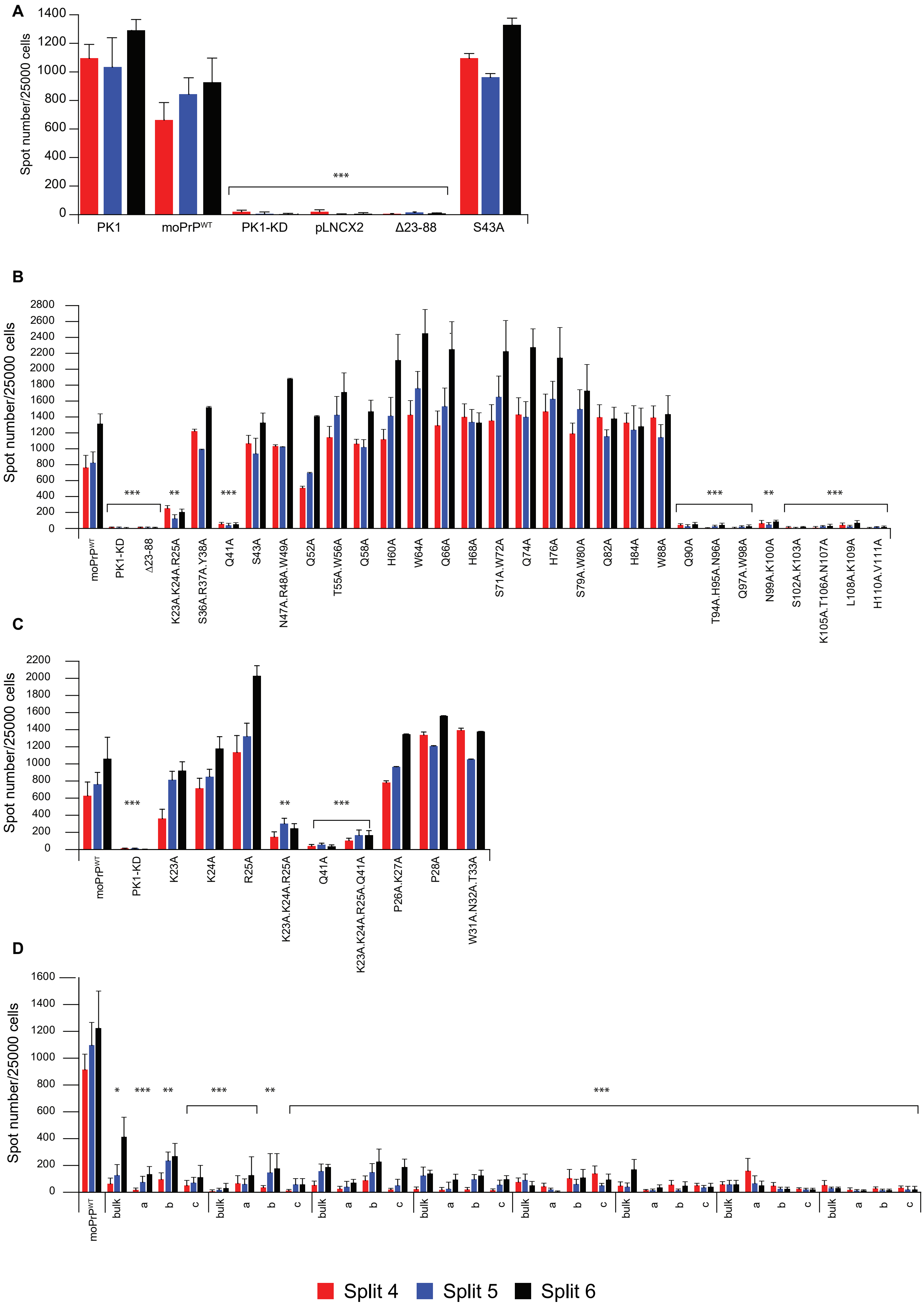


Extended Fig 2

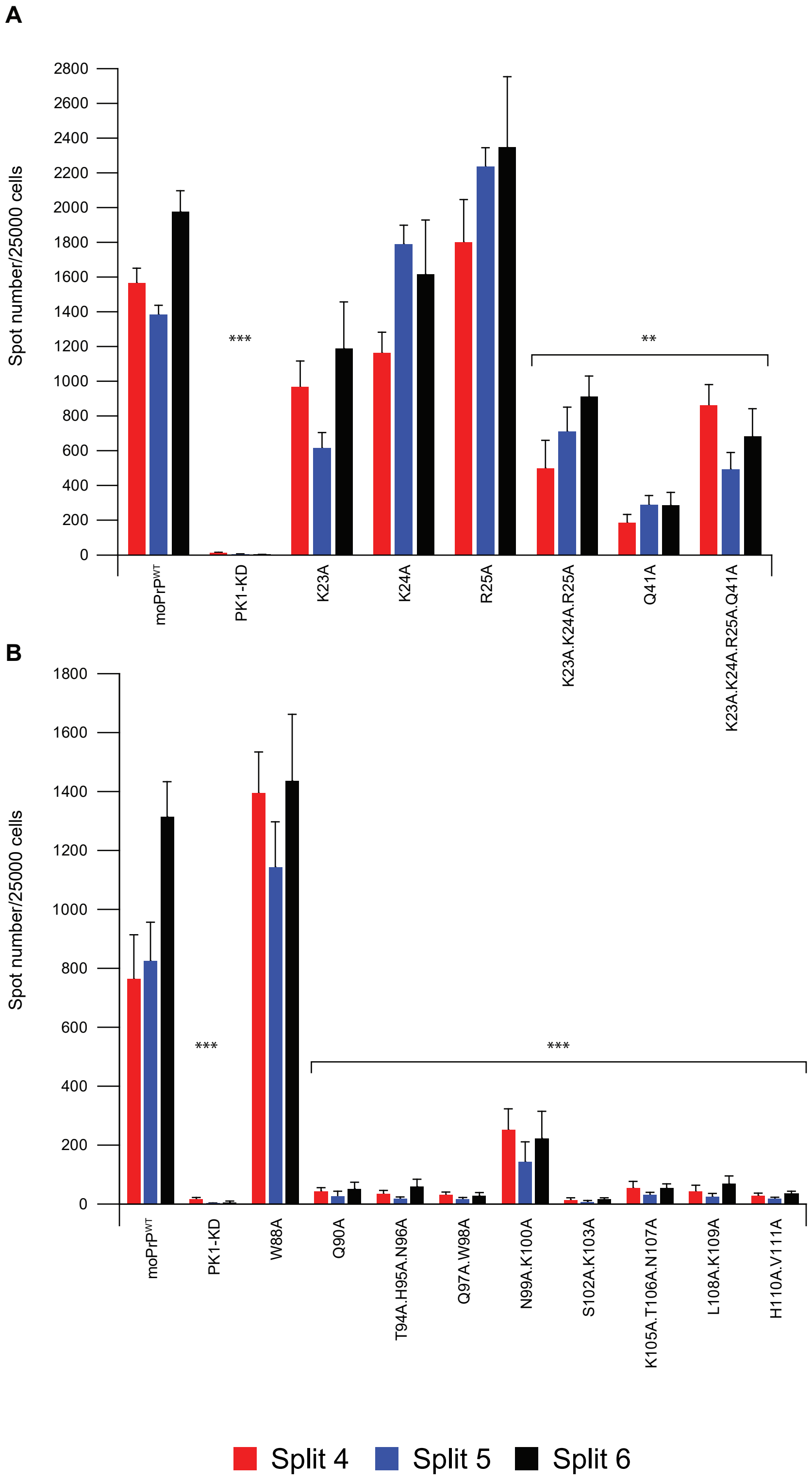


Extended Fig 3

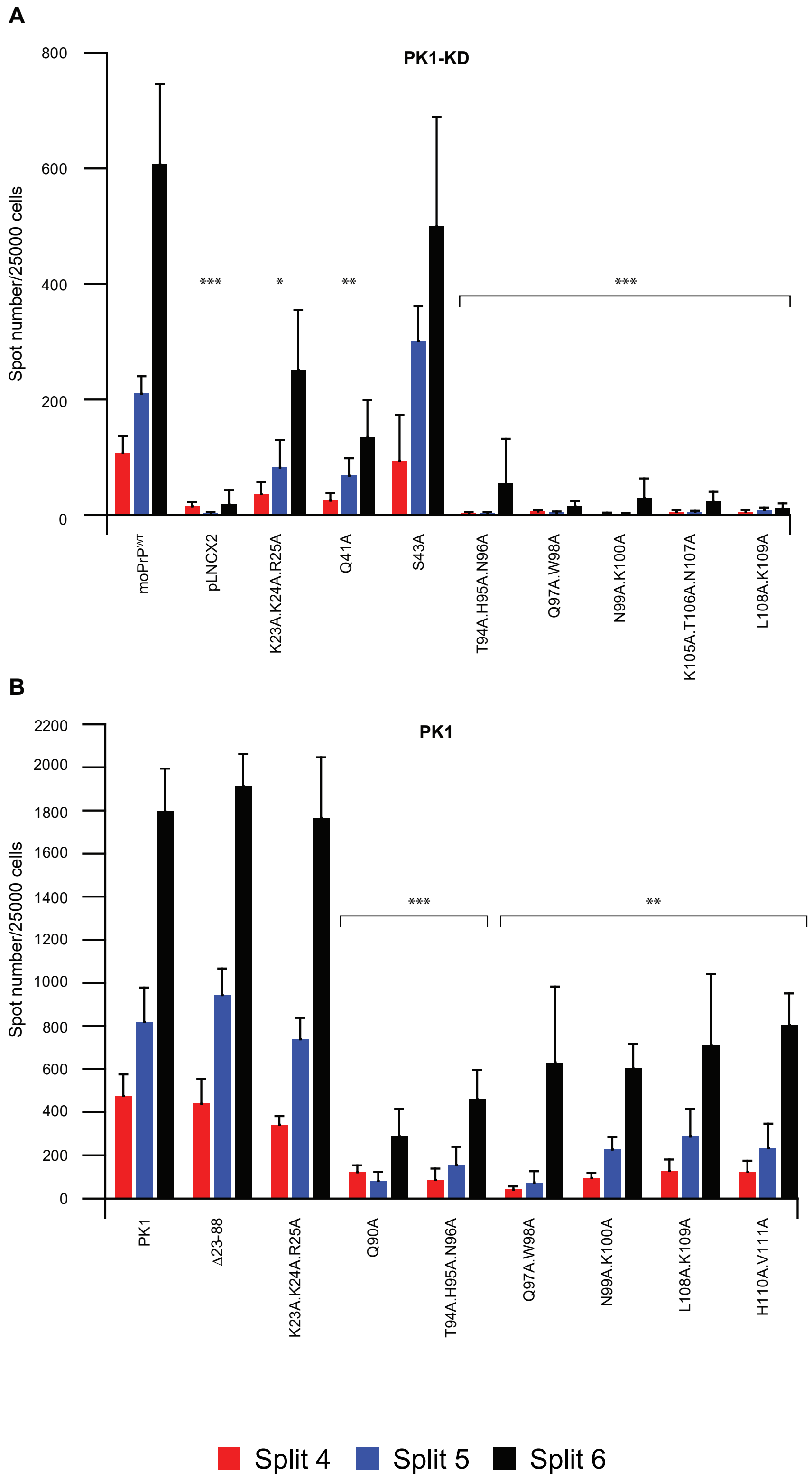


Extended Fig 4

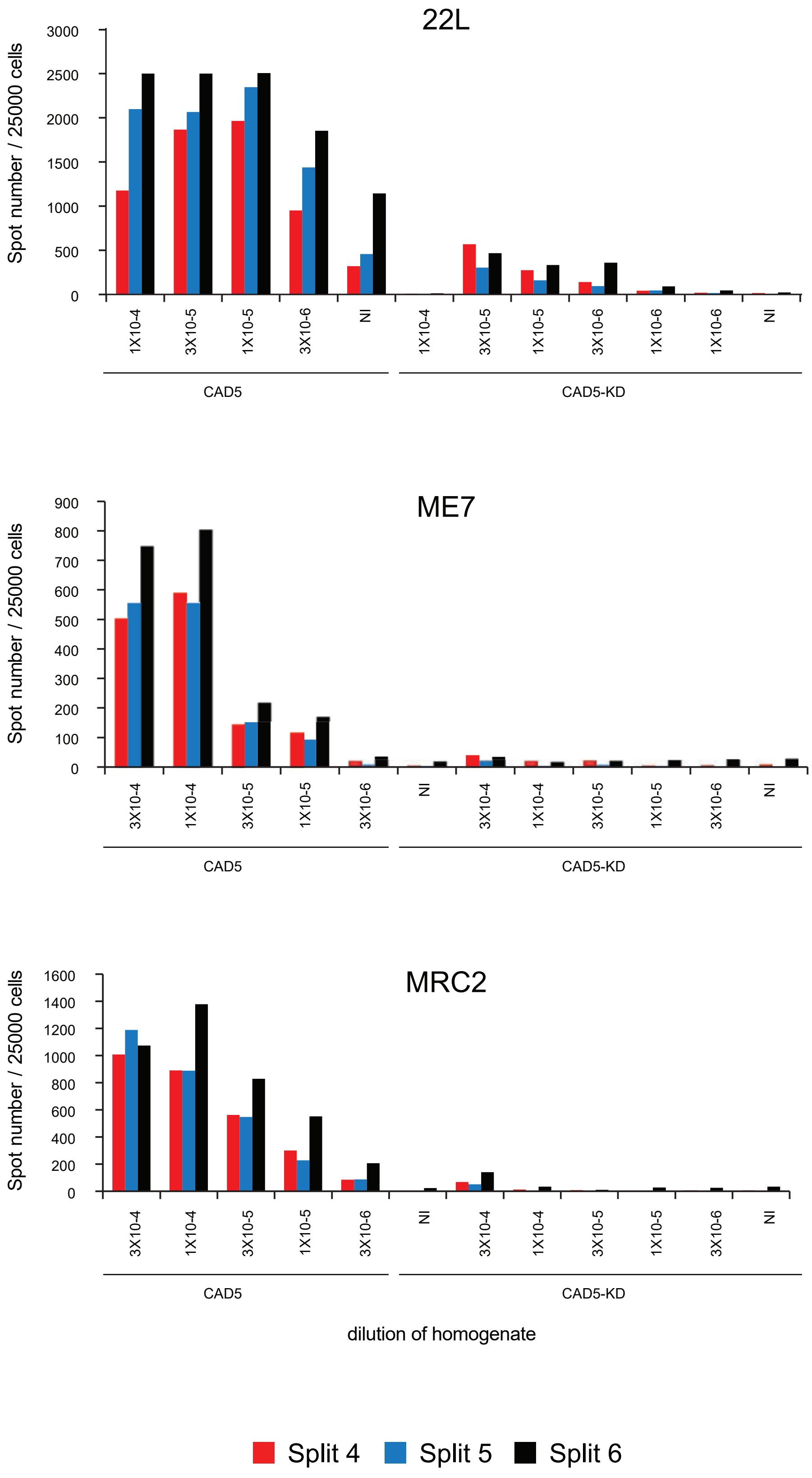


Extended Fig 5

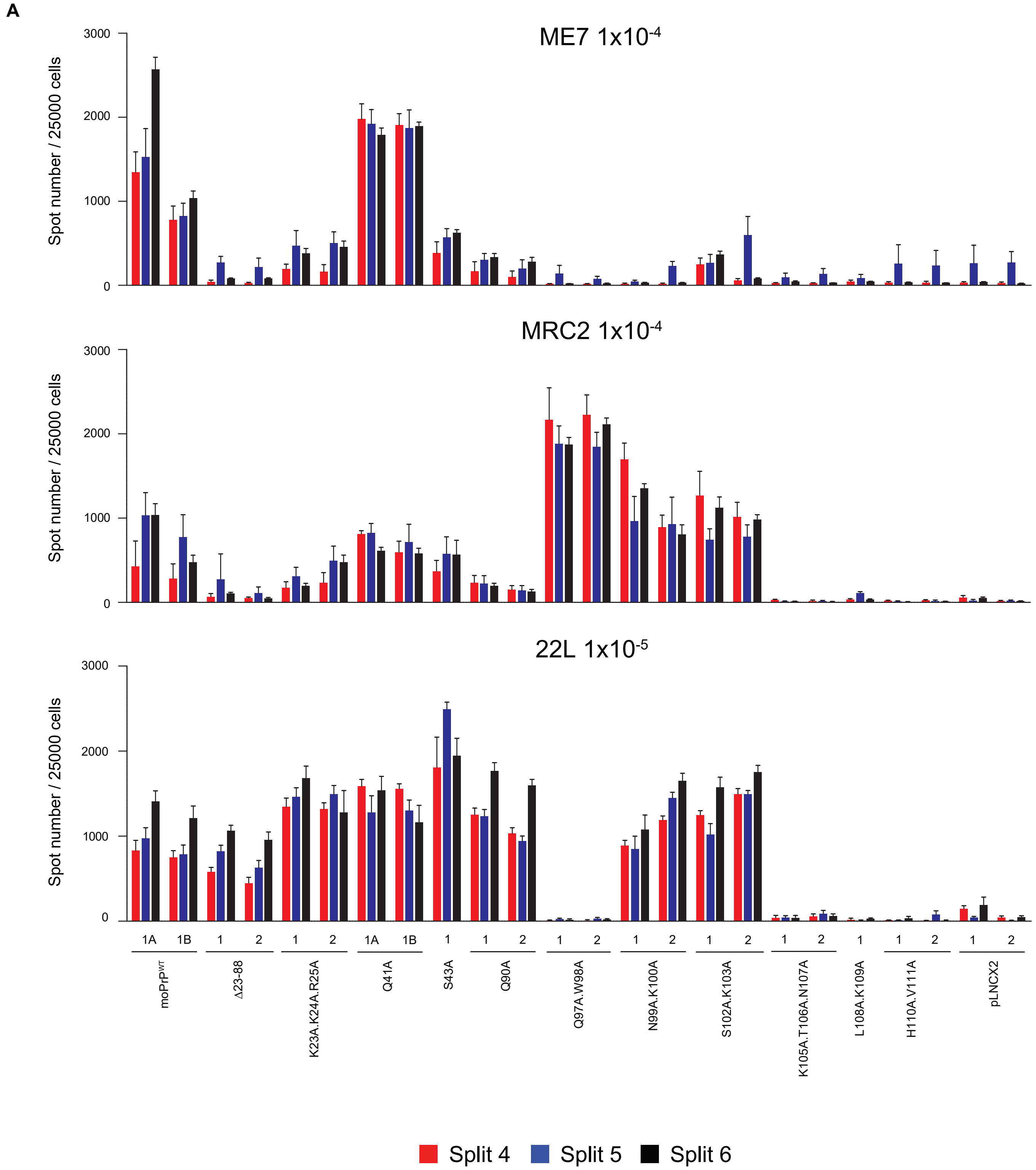


Extended Fig 6

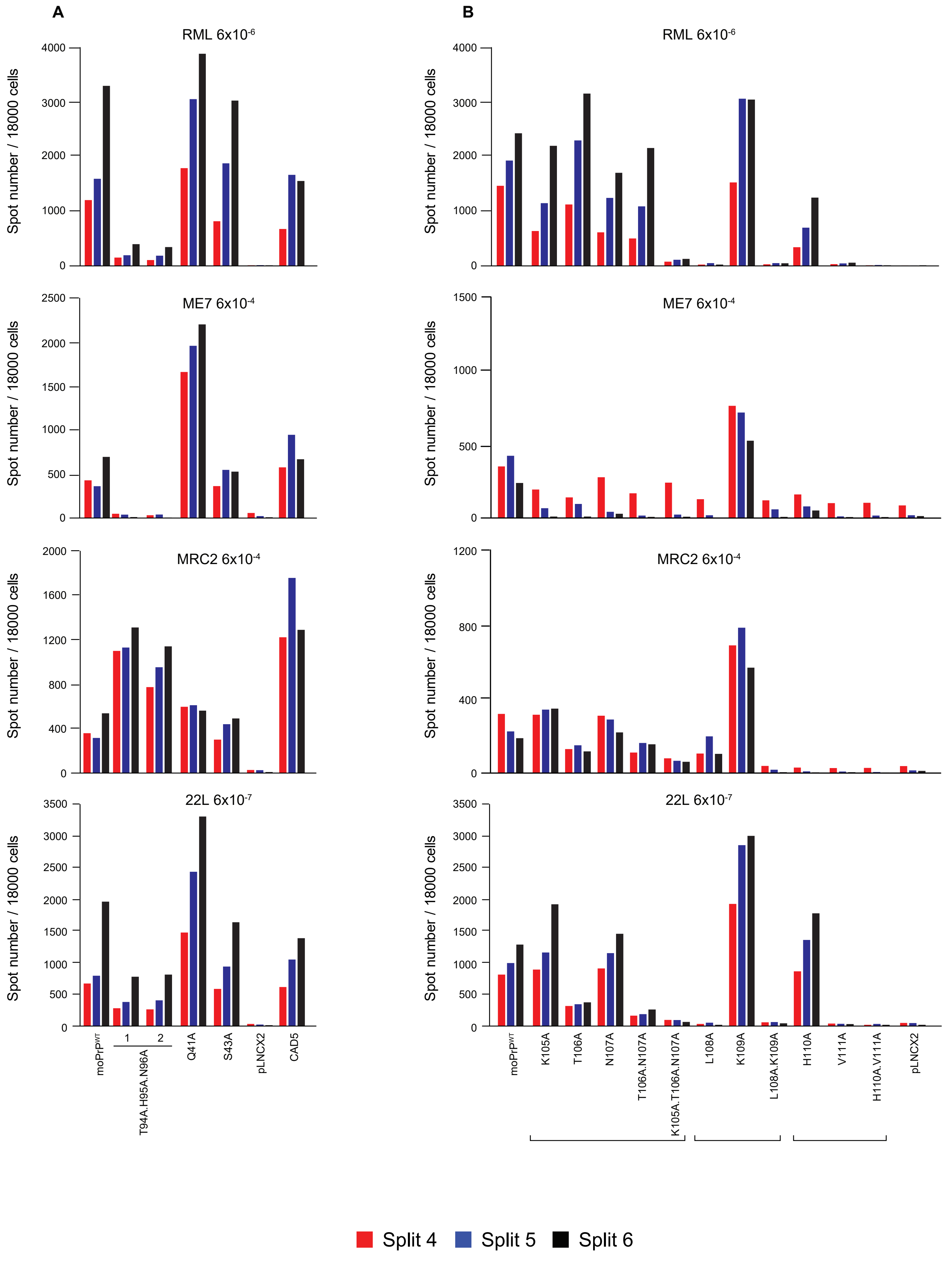


Extended Fig 7

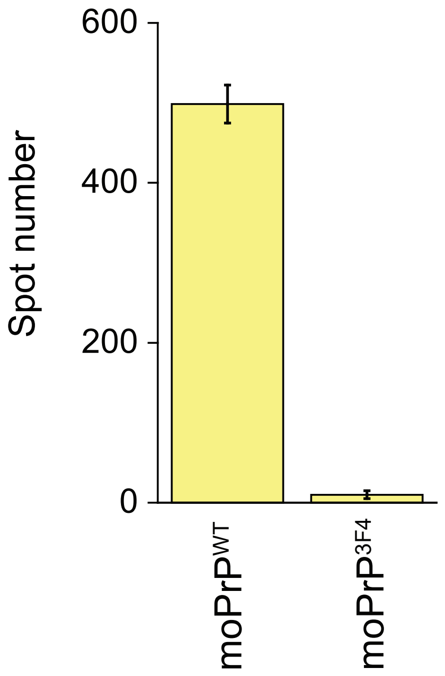


Extended Fig 8
